# Quantifying and mitigating recorder-induced variability in ecological acoustic indices

**DOI:** 10.1101/2023.10.16.562620

**Authors:** David Luna-Naranjo, Juan D. Martínez, Camilo Sánchez-Giraldo, Juan M. Daza, José D. López

## Abstract

Due to the complexity of soundscapes, Ecological Acoustic indices (EAI) are frequently used as metrics to summarize ecologically meaningful information from audio recordings. Recent technological advances have allowed the rapid development of many audio recording devices with significant hardware/firmware variations among brands, whose effects in calculating EAI have not yet be determined. In this work, we show how recordings of the same landscape with different devices effectively hinder reproducibility and produce contradictory results. To address these issues, we propose a preprocessing pipeline to reduce EAI variability resulting from different hardware without altering the target information in the audio. To this end, we tested eight EAI commonly used in soundscape analyses. We targeted three common cases of variability caused by recorder characteristics: sampling frequency, microphone gain variation, and frequency response. We quantified the difference in the probability density functions of each index among recorders according to the Kullback-Leibler divergence. As a result, our approach reduced up to 75% variations among recorders from different brands (AudioMoth and SongMeter) and identified the conditions in which these devices are comparable. In conclusion, we demonstrated that different devices effectively affect EAI and show how these variations can be mitigated.

**Graphical Abstract:** 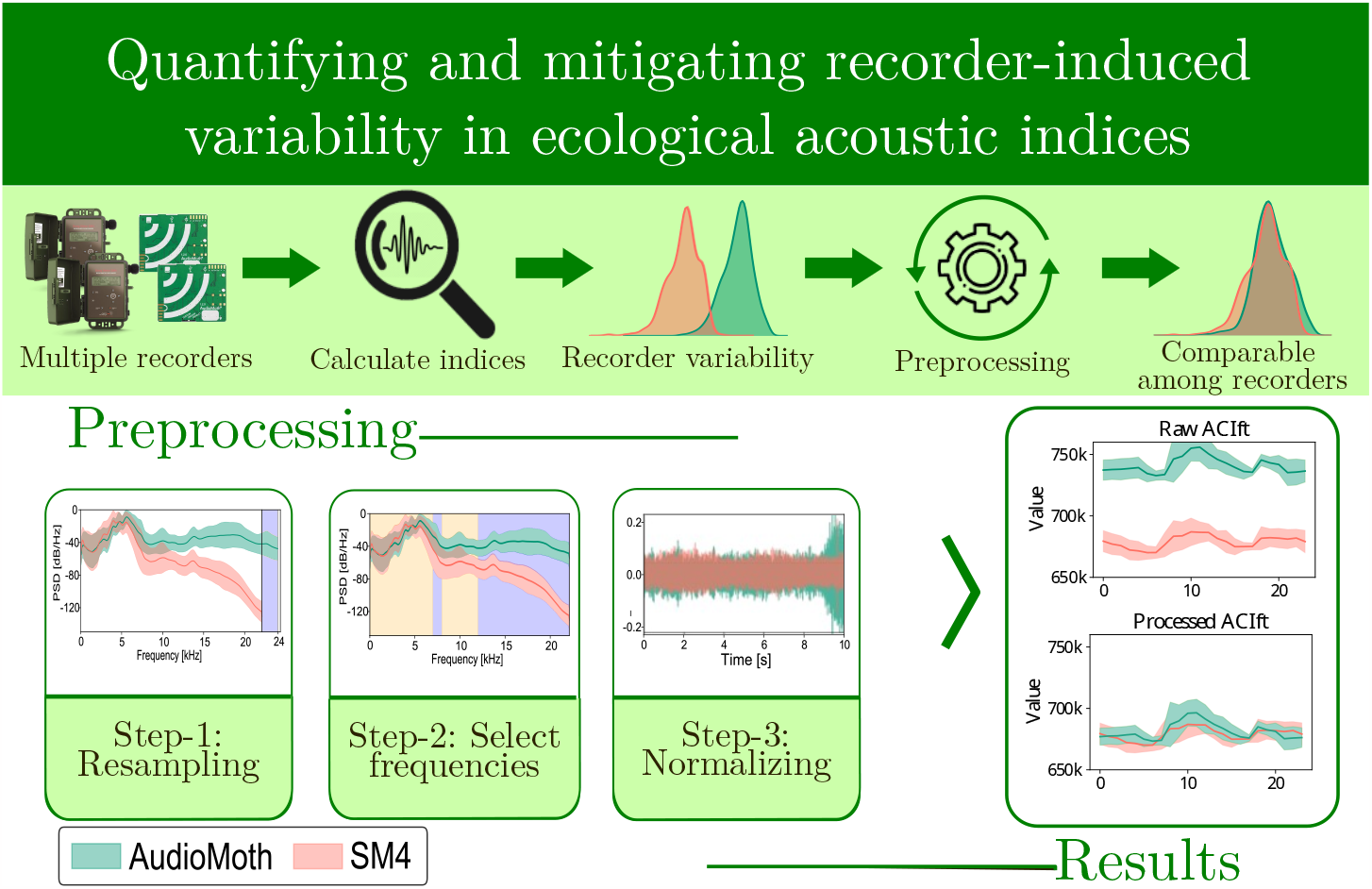

**Highlights:**

- Addressing recorder-induced biases in acoustic indices for improved reproducibility.
- Proposing an effective method to mitigate recorder-related biases.
- Evaluating pipeline proposed performance via acoustic index distribution analysis.

## 1. Introduction

Ecological acoustic indices (EAI) are one of the most common metrics to study soundscapes as they are easy to calculate and are assumed to have biological interpretability (Buxton et al., 2018; Sueur and Farina, 2015; Farina and Gage, 2017). They are essential for large-scale studies using multiple recorders, a growing trend impulsed by the development of new low-cost and commercially available acquisition devices (Gibb et al., 2019) (see (Browning et al., 2017) for a complete list). However, the sensor-dependent sensitivity inherent to using multiple recorders generates a new source of variability that has yet to be studied, as it may affect studies where it is assumed that space and time are the only sources of sound variation. This new source of variability introduces an uncharted realm of potential influence on soundscape studies, underscoring the need to delve into the intricate interplay between sensor characteristics and the resulting acoustic data.

EAI encode complex and biologically relevant information in a single numerical value, efficiently processing large volumes of data (Sueur et al., 2014; Eldridge et al., 2016). These indices are calculated either from the raw signal or in its time-frequency representation to extract different soundscape characteristics such as pitch, occupancy of frequency bands, amplitude, signal gain, differences among frequency bands, and dissimilarity measurements. Due to their flexibility, EAI have been used for different aims, such as acoustic niches, uniformity, soundscape degradation, and environmental studies (Towsey et al., 2014; Shaw et al., 2021; Benoccıet al., 2020). Moreover, EAI are commonly employed as proxies for biodiversity metrics and considered a practical alternative tool for rapid biodiversity assessments Sueur et al. (2008).

However, several studies using the same indices provide contradictory results by correlating with biological variables, generating doubts about EAI effectiveness, despite the prevalence of their use Bradfer-Lawrence et al. (2019a). Consequently, although different works aim to identify the possible causes of this variability and propose alternatives to measure the effectiveness of EAI for characterizing various habitats, a straightforward standard implementation is needed (Ross et al., 2021; Zhao et al., 2019a; Jorge et al., 2018).

The intra-study variability of EAI has been attributed to various factors, such as the sensitivity to changes in environmental conditions, noise, and the sampling protocol Bradfer-Lawrence et al. (2019a); Metcalf et al. (2021a); Lillis and Mooney (2022); Zwerts et al. (2022). The literature review (see Appendix A) highlights ecoacoustic data variability reduction strategies, yet it predominantly focuses on intra-brand evaluations using filtering methods and sampling adjustments for EAI Hyland et al. (2023a). Inter-brand comparisons are notably absent, preventing a comprehensive understanding of variability and emphasizing the need for broader cross-brand research for effective mitigation strategies. This problem hinders reproducibility and a direct comparison of EAI among studies.

Additionally, the current trends in covering large geographic areas with multiple recorders (Gibb et al., 2019) and the design of long-term studies that compare data acquired in different years and with different devices (Abrahams et al., 2023) worsen the issues mentioned above (even with same-brand devices). Furthermore, the increased number of recorders within the same study, sometimes of different brands and characteristics, and the deterioration of the microphones induce a sensor bias that prevents the direct comparison of EAI. This problem will increase since nowadays researchers commonly have in their stocks high and low cost devices (SongMeter and BAR and AudioMoth) (Hill et al., 2019). The use of devices with different characteristics thwarts the development of standardized acoustic monitoring methodologies, curtailing comprehensive ecological insights. In more critical scenarios, this limitation may lead to misguided conclusions about habitat conditions. Consequently, it is essential to reduce the influence of the recording device in ecoacoustic studies by developing methodologies that produce comparable acoustic features (e.g., EAI), independent of the acquisition device.

In this study, we address the challenge presented by this open issue, which mainly affects the growth of acoustic monitoring and is noticeably prominent in large-scale studies (Gibb et al., 2019). The novelty of our work resides in the proposition of a methodology to diminish this sensor-induced variability. Our primary objective is to enable researchers to effectively compare EAI among diverse studies (or sites), even when employing recorders from different brands. Therefore, we analyzed how the use of different recorders (from the same and different brands) impacts the EAI calculation. We focused on the eight EAI most commonly used in biodiversity studies: ACIft, ADI, AEI, M, H, NP, BI, and NDSI (Buxton et al., 2018). To this end, we analyzed their analytical formulation, biological hypothesis, and practical implementation to determine the leading causes of variability produced by different hardware and parameters used for data acquisition. Then, with these sources of variability defined, we proposed a preprocessing methodology to reduce sources of variability and test methodology with data from simultaneous on-field recordings. The codes used in this work are shared in a GitHub repository.

## 2. Materials and methods

### 2.1 Data collection

We conducted two field experiments to evaluate the impact of intra- and inter-brand variability when computing EAI. In the first experiment, audio recordings were simultaneously obtained from different brand recorders at the same site (same tree) and replicated in four humid forest habitats (S1 to S4). The second experiment involved deploying a setup of 15 AudioMoth recorders (shown in Figure 1) to assess the impact of using different recorders of the same brand (S5). Geographical and temporal details of these experiments can be found in Table 1.

**Table 1:** Geographical details and configuration parameters of the recorders used for data acquisition to evaluate intra-brand and Inter-brand variability in calculating ecological acoustic indices (EAI).

| Site | Experimental Setup | Recorder Specifications | Locality Data |
| --- | --- | --- | --- |
| S1 | Inter-brand<br>Devices: 2<br>Recordings: 899<br>Sample Scheme: 1 x 20 min | *SongMeter*:<br>- FS 44.1 kHz<br>- Gain: 16dB<br>*AudioMoth*:<br>- FS: 48 kHz<br>- Gain: medium | *Location*: Amalfi, Antioquia-Colombia<br>*Geographic Coordinates*: (-75.16991, 6.83370)<br>*Date*: June 3 to July 13, 2019 |
| S2 | Inter-brand<br>Devices: 2<br>Recordings: 868<br>Sample Scheme: 1 x 20 min |  | *Location*: Anorí, Antioquia-Colombia<br>*Geographic Coordinates*: (-75.09621, 6.96325)<br>*Date*: March 20 to March 30, 2019 |
| S3 | Inter-brand<br>Devices: 2<br>Recordings: 1711<br>Sample Scheme: 1 x 15 min | *SongMeter*:<br>- FS: 48 kHz<br>- Gain: 16dB<br>*AudioMoth*:<br>- FS: 48 kHz<br>- Gain: medium | *Location*: San Rafael, Antioquia-Colombia<br>*Geographic Coordinates*: (-75.03207, 6.26504)<br>*Date*: October 17 to November 04, 2021 |
| S4 | Inter-brand<br>Devices: 2<br>Recordings: 1988<br>Sample Scheme: 1 x 15 min |  | *Location*: San Rafael, Antioquia-Colombia<br>*Geographic Coordinates*: (-75.01989, 6.26283)<br>*Date*: November 04 to November 25, 2021 |
| S5 | Intra-brand<br>Devices: 15<br>Recordings: 203<br>Sample Scheme: 1 x 5 min | *AudioMoth*:<br>- FS: 48 kHz<br>- Gain: medium | *Location*: San Miguel, Antioquia-Colombia<br>*Geographic Coordinates*: (-75.596635, 6.022450)<br>*Date*: July 24 to July 25, 2022 |

**Figure 1.**
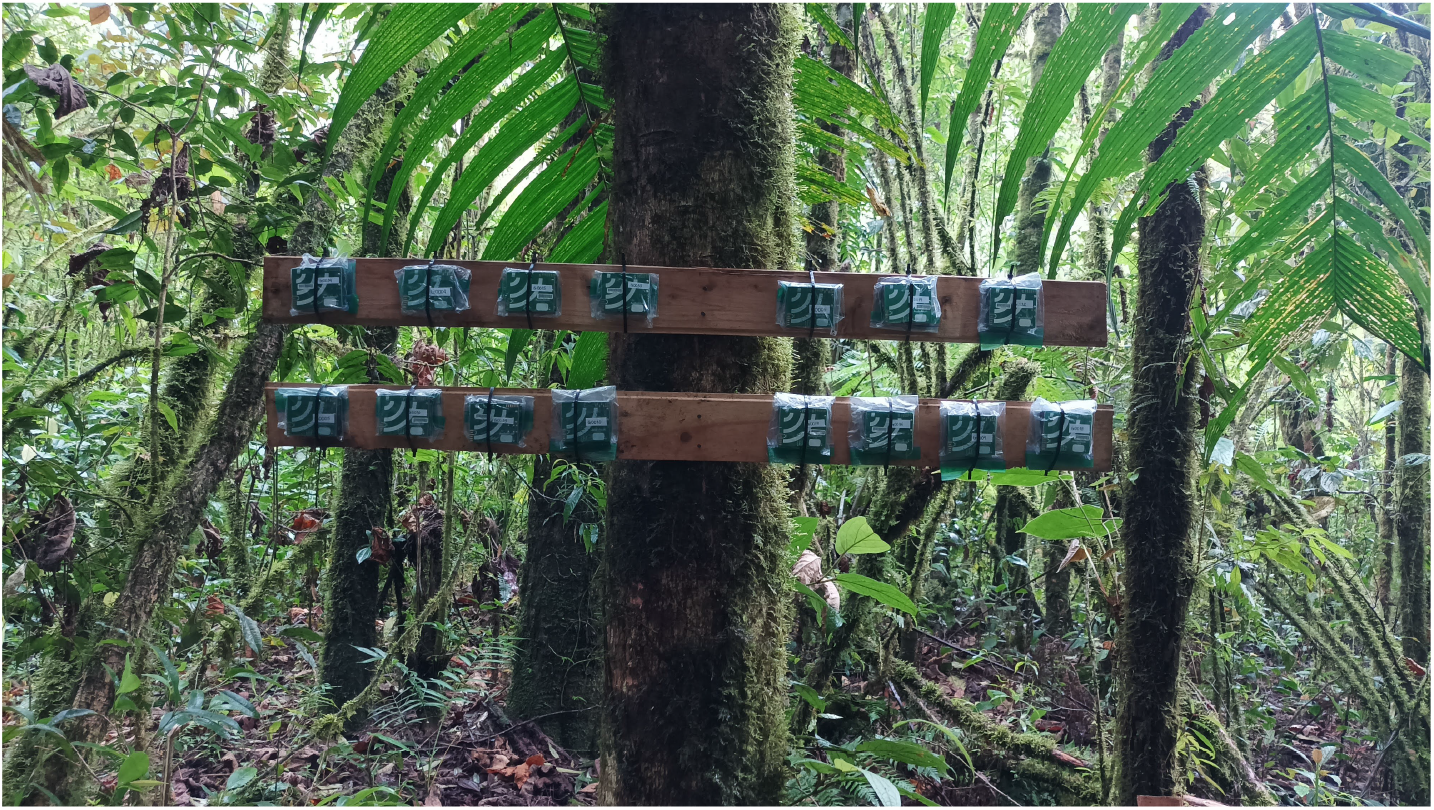
Fifteen recorders (five each of AudioMoth v1.0, v1.1, and v1.2) were deployed for evaluating the impact of single brand device variations (San Miguel, Antioquia, Colombia).

The recordings were made with two of the most popular references nowadays. The first one is the SongMeter 4 (SM4) recorder from Wildlife Acoustics (www.wildlifeacoustics.com), which is considered a high-quality data acquisition device (Browning et al., 2017). It has 16-bit programmable stereo recording, a maximum sampling frequency of FS=96kHz, and two omnidirectional microphones with a sensitivity of −35 ± 4 dB as specified in its datasheet (acoustics Bioacoustics Monitoring Systems). The second is the AudioMoth recorder (www.openacousticdevices.info), which has gained enormous popularity in recent years as a low-cost (approximately USD90) open-source device. The AudioMoth is a mono recorder; it can record up to 16 bits and is designed to capture audible and ultrasonic sounds with a maximum frequency sample of FS=384kHz. It has a built-in MEMS (Micro Electrical-Mechanical System) microphone.

Sites S1 and S2 were equipped with two devices: a SongMeter 4 (SM4) and an AudioMoth (version 1.0.0), both configured according to the technical parameters specified in Table 1. At sites S3 and S4, the SM4 frequency sample rate was adjusted to 48 kHz. Site S5 had fifteen AudioMoths: five V1.0, five V1.1, and five V1.2. These devices were systematically installed in an interleaved manner within the designated setup (se Figure 1). All devices were identically configured, following the settings presented in Table 1.

### 2.2. Analytical EAI analysis

For this study, we selected the eight EAI most commonly used in ecoacoustics, as evidenced by Buxton et al. (2018) and Alcocer et al. (2022). These indices quantify acoustic variability at the sampling sites, provide different information about the soundscape, and their use spans different habitats and environmental scales Zhao et al. (2019b). In addition, their mathematical methods are the basis on which many other EAI are defined. We then analyzed the theory behind their analytical formulation and identified their advantages and leading causes of sensitivity. The complete analysis is included in Appendix B and is summarized in Table 2.

**Table 2:** Overview of the eight most commonly used EAI: advantages, disadvantages, and sensitivity to sources of variability.

| Description | Advantages | Sensitivity |
| --- | --- | --- |
| Acoustic complexity index (ACIf<sub>t</sub>)<br>It quantifies the acoustic complexity based on the intrisectral variability of the frequency bands (see Appendix B.1) | It is robust to using multiple recorders with different gains (e.g., due to different brands of microphones) | - Different sampling frequencies<br>- Geophonic sources (such as rain or wind)<br>- Non-constant sources of anthropophony |
| Acoustic diversity index (ADI)<br>It determines the occupancy degree of acoustic niches (see Appendix B.2) | It is robust against sampling frequency variations because the frequency domain is collapsed. | - Binarization threshold for the histograms (microphones with different gains).<br>- Frequency response of the microphones<br>- Artifacts with energy content in several frequency bands, such as rain. |
| Acoustic evenness index (AEI)<br>It estimates the uniformity of species in different acoustic niches (see Appendix B.3) | Same as ADI | Same as ADI |
| Median of the amplitude (M)<br>It uses the mean amplitude envelope as an indicator of sound sources in the habitat (see Appendix B.4) | It encodes the amplitude of the recording | - Microphone distance and the habitat structure<br>- Environments with high-intensity noise sources<br>- Microphones with different gains |
| Acoustic Entropy Index (H)<br>Indicator of heterogeneity based on Shannon uniformity over time and frequency (see Appendix B.5) | It reduces its sensitivity to gain differences among devices and different sampling frequencies. | Broadband noise sources (such as traffic, rain, or wind) |
| Number of Peaks index (NP)<br>Indicator of acoustic complexity using the number of prominent peaks in frequency as a spectral profile of the acoustic richness of the habitat (see Appendix B.6) | - It is less sensitive to environment noise<br>- It is robust to different gains and sampling frequency differences among devices | Frequency response of the microphones |
| Bioacoustic index (BI)<br>It evaluates the relative abundance of species in the frequency bands of the biophony (see Appendix B.7) | It is less sensitive to microphone gain | Frequency response of the microphones |
| Normalized Difference Soundscape Index (NDSI)<br>It estimates the level of anthropogenic disturbance in the soundscape (see Appendix B.8) | Same as BI | Same as BI |

This theoretical analysis shows that the three issues (related to using different recorders) that most affect these indices (and are commonly ignored or mistreated by researchers) are:

1. Audios with different sampling frequencies (FS): The maximum frequency of interest is critical to select the FS with different devices. Many EAI are based on the Fast Fourier Transform (FFT) of the signal. The resolution of this representation is a ratio between FS and the number of samples (nfft) in the window segment. The size of the frequency bins (partitions) is given by FS/nfft. If two recordings have different FS, the indices will be biased.

2. Gain variations at different frequencies: The frequency response of the microphones indicates how the input vs. output varies at different frequencies. This behavior is commonly sought to be linear; however, this is uncommon in recorders used for passive acoustic monitoring. Microphones of different characteristics will have a different dynamic response, especially at higher frequencies. For example, according to its datasheet (acoustics Bioacoustics Monitoring Systems), the SongMeter 4 microphone becomes non-linear after 8kHz.

3. Differences gains among recording devices: The signal amplitude of a microphone depends on the sound pressure that the microphone may capture. This depends on the manufacturing materials and its level of degradation. Significant differences in gain are expected from microphones of different brands or after severe outdoor use.

### 2.3. Proposed preprocessing approach

Fig. 2 shows the proposed preprocessing methodology to tackle the above-mentioned issues. It consists of three steps:

**Figure 2.**
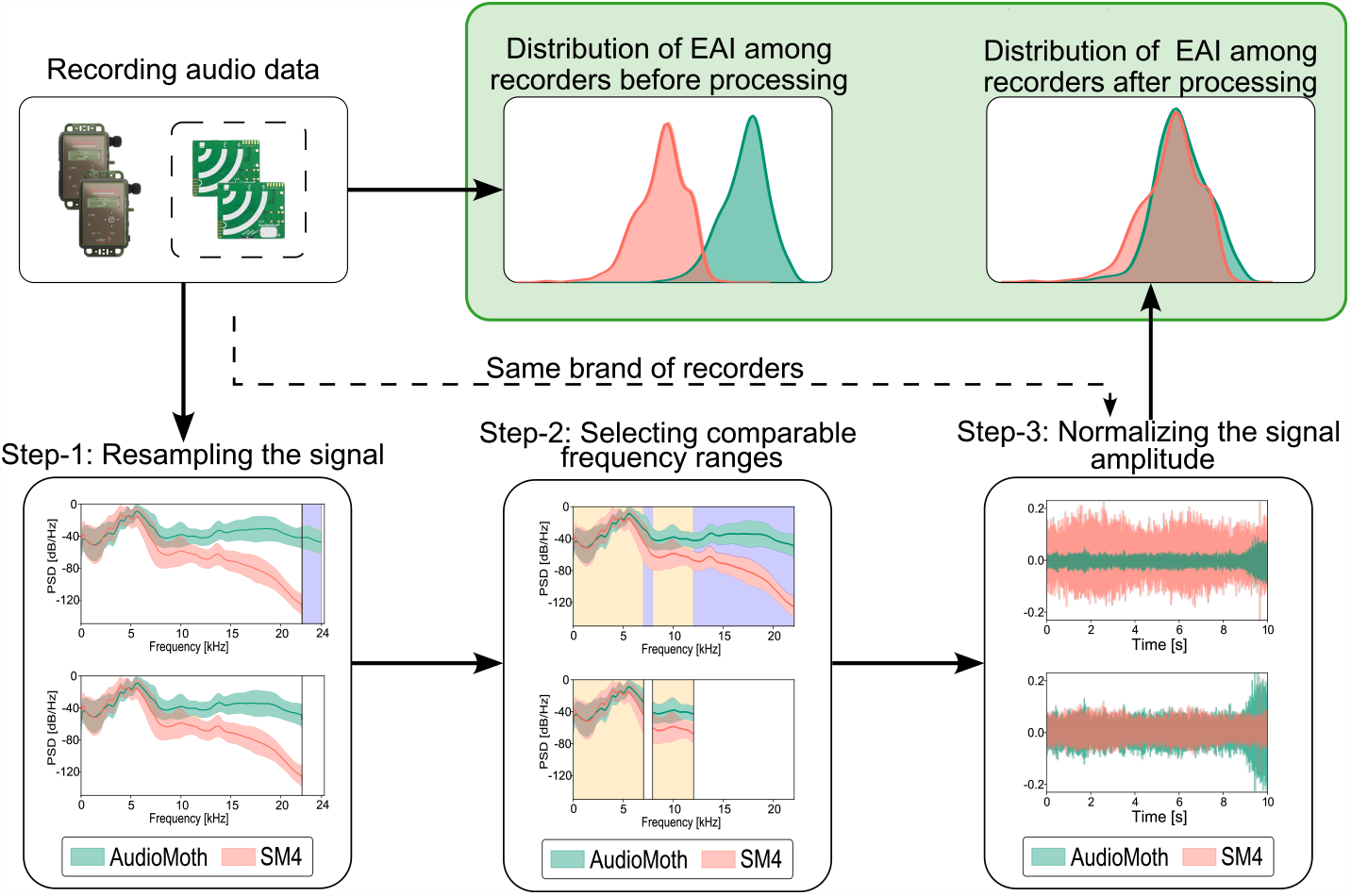
Proposed pipeline to make EAI comparable across different recorders. Purple bands indicate intervals that are not comparable in frequency due to the maximum frequency in Step-1 and the frequency response in Step-2. The yellow bands indicate comparable frequency intervals in Step-2. For recorders of the same brand and FS, only Step-3 is required.

Step-1 Resampling the signal: If audio recordings have a different FS, they must be subsampled to the lowest one. Otherwise, extrapolation would be required, and it would introduce artifacts. We recommend subsampling with the Python PyTorch library, which has a resampling function based on the Kaiser windows Liu et al. (2020). This method was selected because of its low attenuation of high frequencies and low computational burden, an essential criterion for processing the large volumes of data handled in ecoacoustics (the computational burden can grow exponentially with some methods).

Step-2 Selecting comparable frequency ranges: Start averaging the Power Spectral Density (PSD) of all recordings from each device and then use coherence to determine the frequency range in which different devices have similar frequency behavior. Only within this frequency range the indices are comparable among devices. Coherence measures the linear frequency dependency between two signals with the same bandwidth and is commonly used to assess the similarity between different signals. Coherence between signals *x* and *y* is defined as:

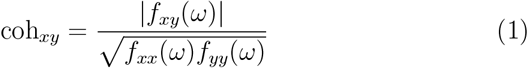

where *f*_*xy*_(*ω*) is the average cross-covariance of *x* and *y*, also known as the cross-spectral density, with *ω* as frequency components of signals. *f*_*xx*_(*ω*) and *f*_*yy*_(*ω*) correspond to the average spectral density (or auto-correlation) functions of *x* and *y*, respectively. Coherence is a symmetric and bounded measure between 0 and 1, where a value of |*coh*_*xy*_(*ω*)| = 0 indicates that the signals have a perfect linear relationship in frequency and a |*coh*_*xy*_(*ω*) | = 1 indicates that there is no linear relationship between the signals in frequency.

Calculate the average coherence over a sliding frequency window with a bandwidth of 1 kHz (the same partition size used for EAI such as Acoustic diversity index ADI and Acoustic evenness index (AEI)) to compare recordings using coherence. The median coherence can then determine common patterns across recordings and the comparable frequency ranges among devices. Moreover, this approach facilitates the comparison of acoustic properties.

Step-3 Normalizing the signal amplitude: Normalize each audio with its root mean square (RMS) value as follows:

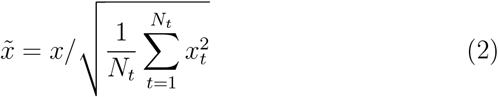

where 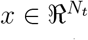 is a mono signal. Normalizing each recording by its RMS value guarantees that we are not altering the heterogeneity information in the audio recording. With this, signals from all devices will be in the same range of scales, reducing their impact on the signal amplitude. A common mistake is to normalize all recordings to a specific amplitude. It will cause the temporal indices to be biased to that value, making it impossible to compare, for example, different sites in a landscape.

The proposed methodology can be simplified if all recordings come from same-brand devices and have the same FS. The only issue to account for is variations in gain (Step-3) due to wear and tear of the recorders or manufacturing issues. In this study, all indices were calculated using the Python scikit-maad library proposed by Ulloa et al. (2021). To facilitate reproducibility, all the necessary codes to replicate the proposed pipeline are available in the following GitHub repository: https://github.com/davidluna-fn/recorder-variability-EAI.git

### 2.4 Data analysis

To validate our approach, we compute the eight EAI’s probability density functions (PDF) at each site using the Parzen method Robert (1976). We then compare them between recorders and analyze the results in the three issues related to using different recorders: audios with different sampling frequencies, gain differences among recording devices, and gain variations at different frequencies, to determine how much the PDF varies among devices due to these issues. Then, we applied the proposed pipeline and repeated the comparison to determine if the variability is effectively reduced.

We evaluated the effectiveness of the proposed pipeline by using two metrics: the Kullback-Leibler divergence (KLD) and the Kolmogorov-Smirnov distance (K-S distance). The KLD measures the difference between two PDFs, quantifying the reduction in variability between devices before and after applying the pre-processing steps. We used the symmetric version of KLD (Van Erven and Harremos, 2014), which is unbounded and provides a measure of the variation magnitude. On the other hand, the K-S distance measures the maximum difference between the cumulative probability distribution functions of two samples. This metric is bounded between 0 and 1, where a value of 0 indicates that the distributions are identical, and a value of 1 indicates that they are entirely different. The advantage of using K-S distance is that it is sensitive to both the shape and position of the probability distribution, providing a clear comparison between distributions (James, 2006).

## 3. Results

### 3.1 EAI comparison from raw data

Figure 3 displays the PDFs of the EAI calculated from the recordings of site S1 before applying our proposed pipeline (sites S2, S3, and S4 are presented in Appendix C, Appendix D, and Appendix E, respectively). All PDF discrepancies coincide with the theoretical analysis of Table 2. The FS difference between devices (44.1kHz vs. 48kHz) strongly affected the mean of ACIft and M in S1 and S2, but S3 and S4 did not show this difference since they were recorded with the same FS.

**Figure 3.**
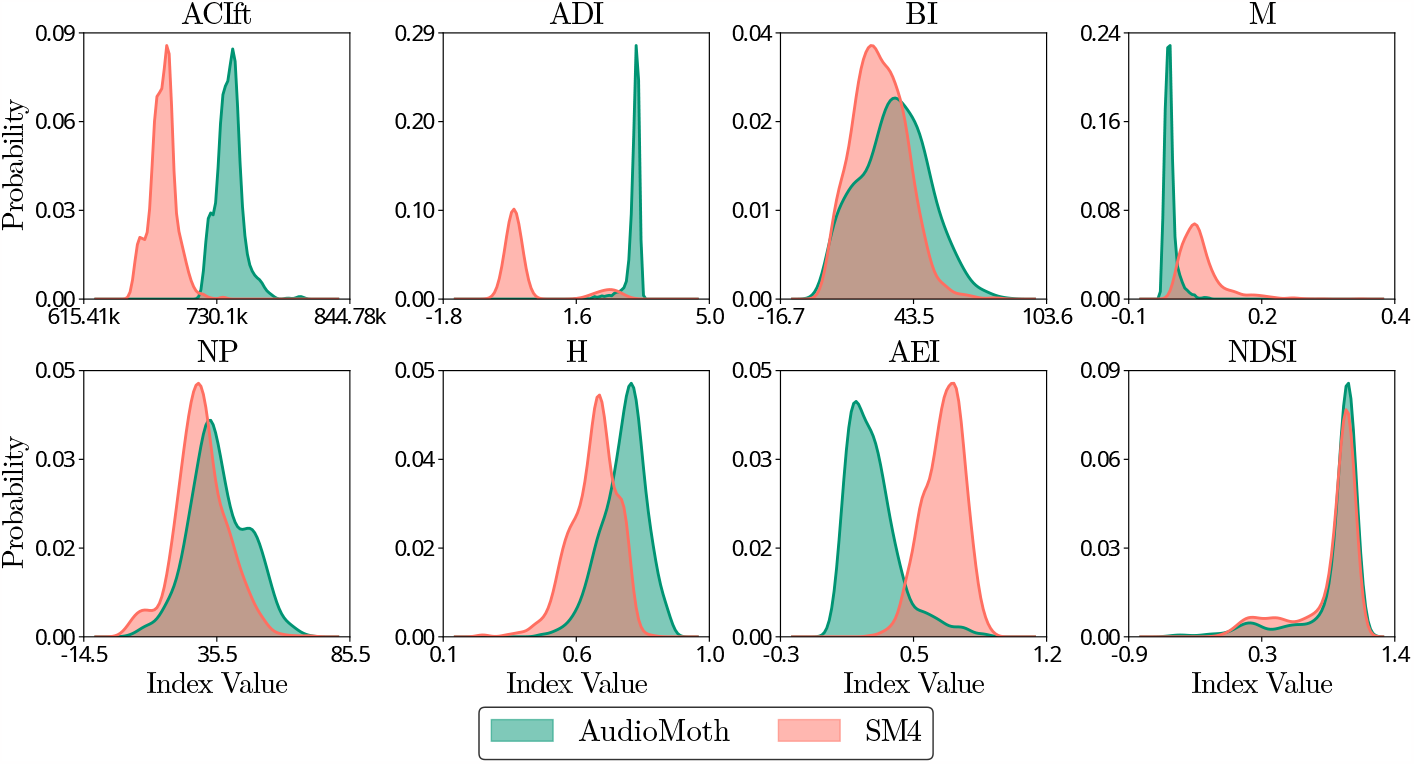
PDFs of eight raw EAI calculated using SM4 and AudioMoth recorders for site S1.

Furthermore, the frequency response impacted the ADI, AEI, H, and NP indices. The PDFs of these indices show different shapes and scales between devices, and this behavior was consistent across all sites. Additionally, the PDFs of M and H varied due to different gains among devices. Finally, we did not identify differences among recorders of the same brand across sites(i.e., S1 and S2 vs. S3 and S4). Next, we will present the results of applying each step of the proposed methodology to reduce the three sources of variability.

### 3.2 Applying the preprocessing methodology

In Step-1, the FS of the AudioMoth device (48kHz) was reduced to FS=44.1kHz after applying the sub-sampling proposed. In Step-2, the frequency response of the two devices per site was analyzed using the coherence measure with the proposed 1kHz sliding frequency window to determine the comparable ranges between the two brands. Figure 4 presents the results for site S1, which show that the frequency response of the recorders is comparable up to 12kHz, where the SM4 falls below the threshold. This behavior is consistent for all studied sites.

**Figure 4.**
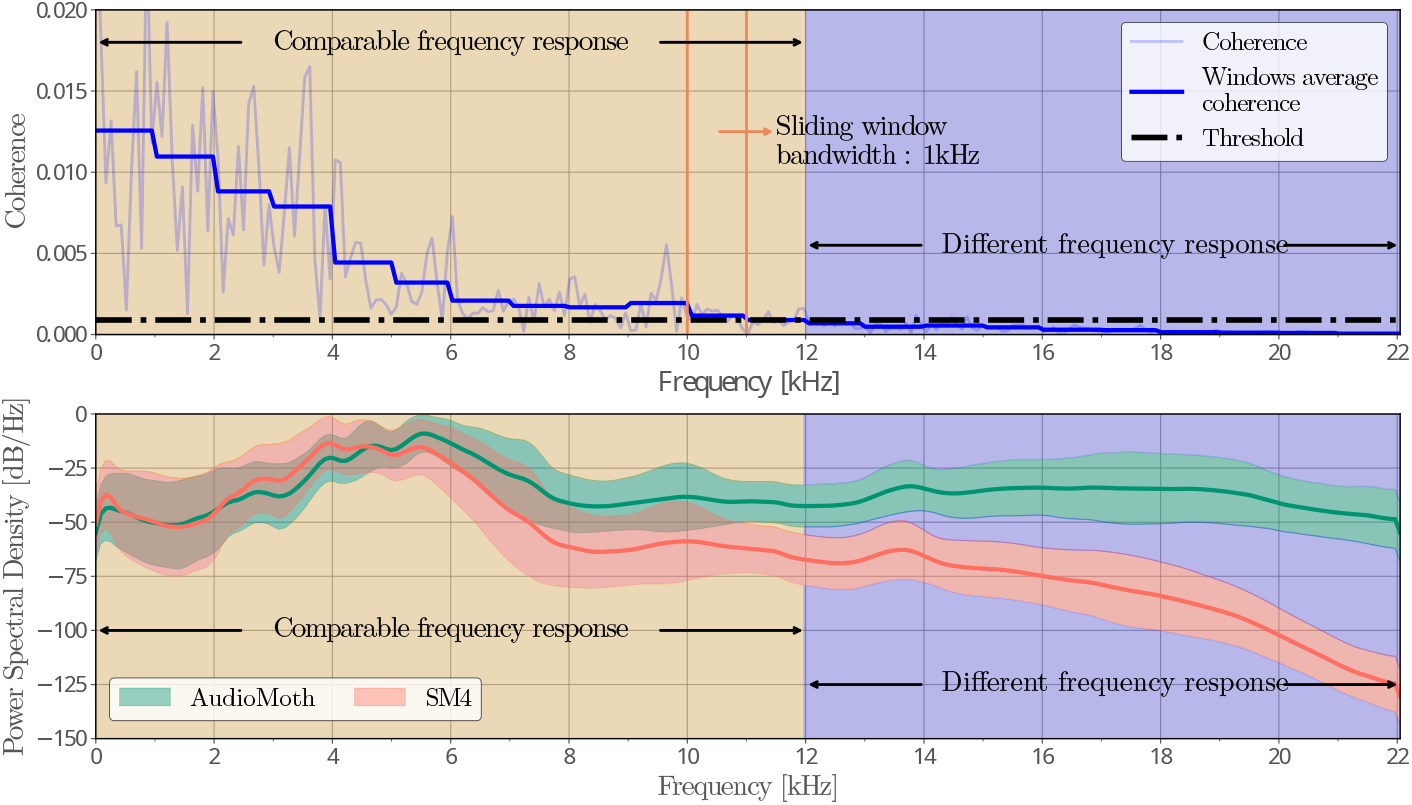
(Top) Coherence sliding window for detecting comparable frequency ranges between recorders. (Bottom) Power spectral density average of subsampled signals.

The coherence measure results coincide with what is visually evident in the frequency responses of the recorders. After 12 kHz, the two devices show different trends, with the SM4 attenuating in high frequencies, consistent with its datasheet, and the AudioMoth maintaining a quasi-linear gain. The bottom panel of Figure 4 illustrates this behavior.

Next, Figure 5 shows the normalization results of Step-3. The top plots display the time series of the same time window recorded by the two devices. The black line represents the median envelope, where the SM4 has approximately four times higher amplitude than the AudioMoth. However, when normalizing the signal using the RMS value, as seen in the bottom plots, the difference in amplitude is eliminated, and the median amplitude is the same for both recorders.

**Figure 5.**
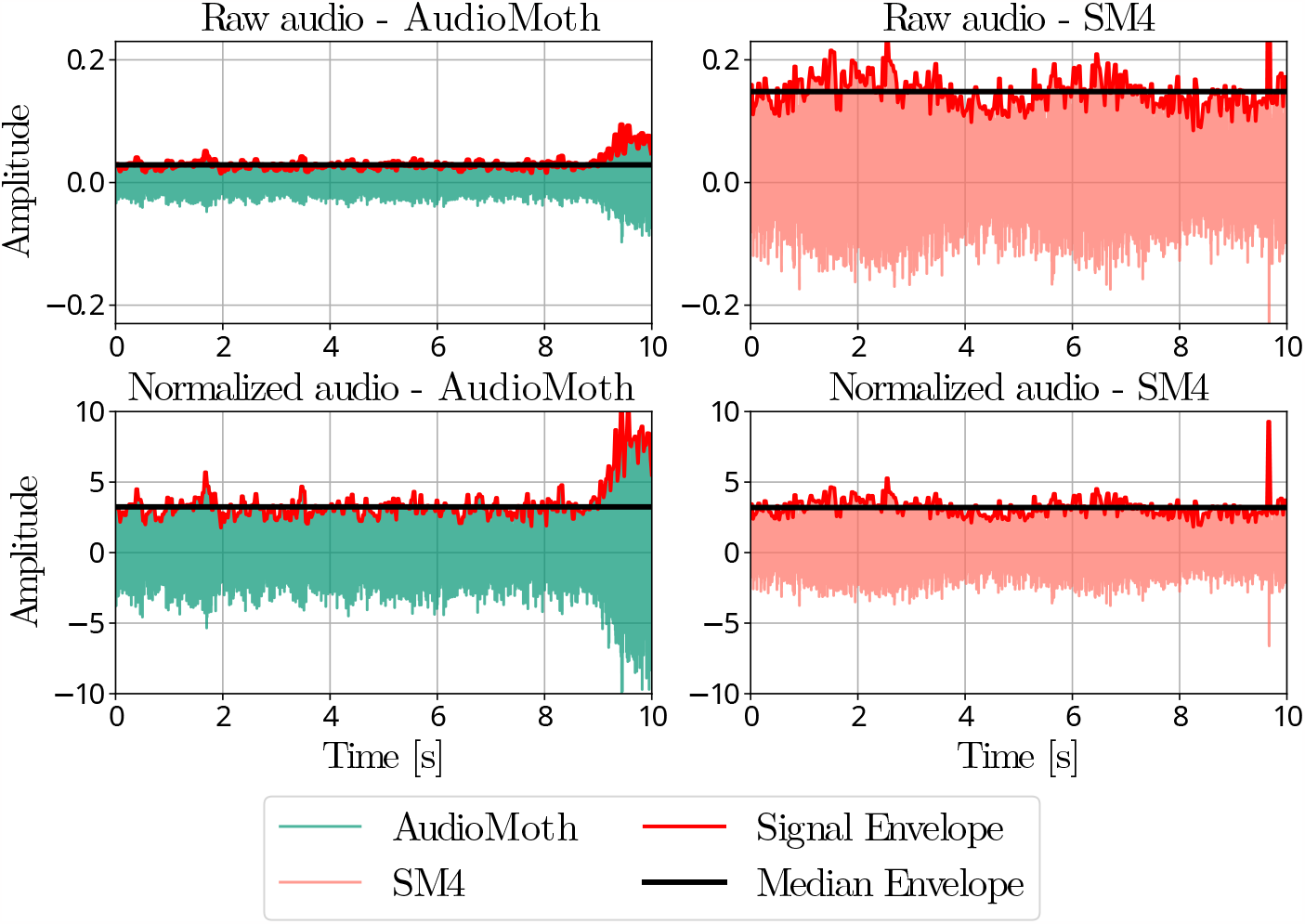
(Top) Amplitude difference between the envelopes of audio signals acquired with different recorders in a 10 s segment. (Bottom) Correction with the proposed normalization.

Finally, Figure 6 shows the mean amplitude of 15 AudioMoth simultaneously recording at S5 before (top) and after (bottom) applying Step-3. After preprocessing, the quantiles represented by the boundaries of the boxplots are closer to the mean, indicating a narrower spread of values.

**Figure 6.**
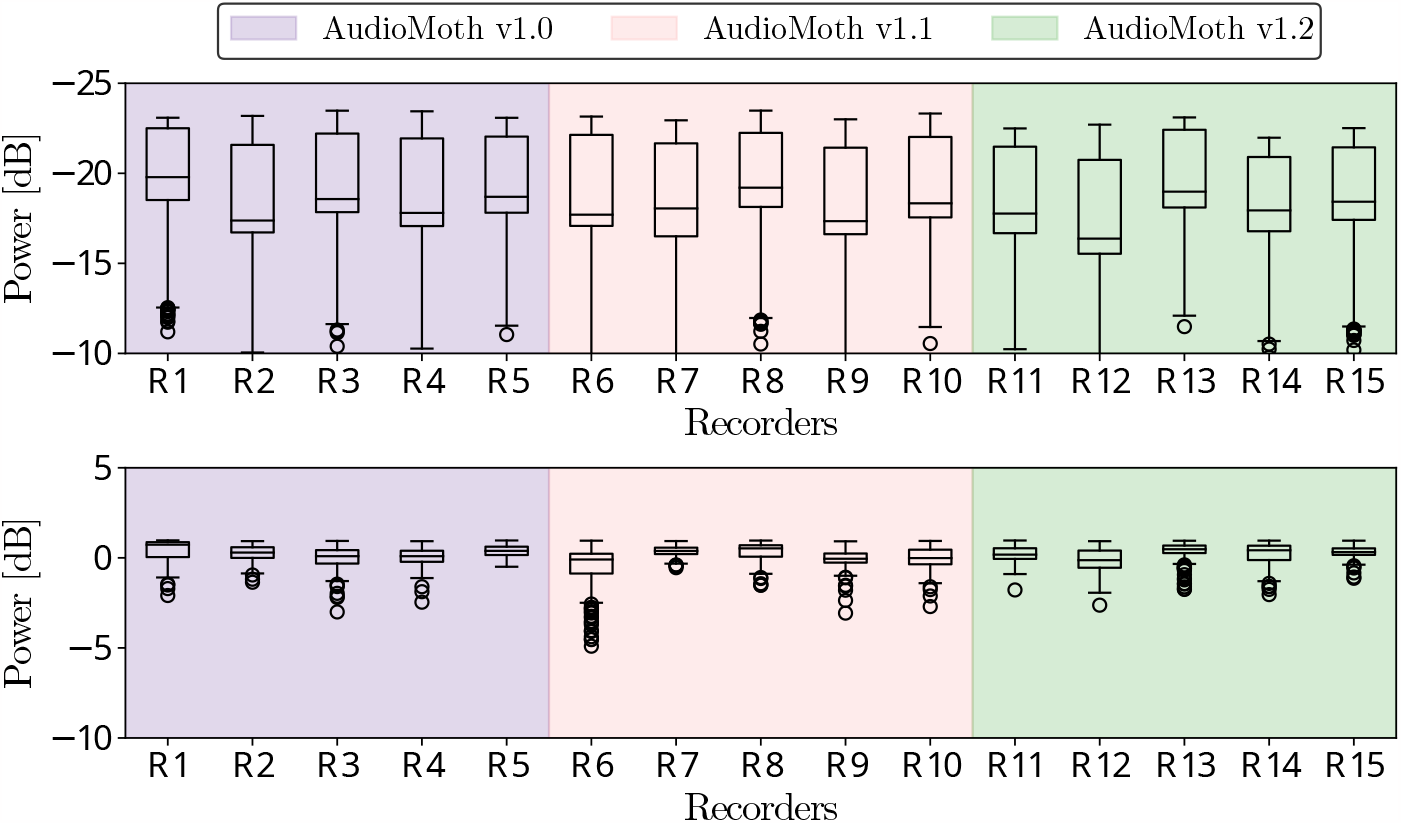
Comparison of mean amplitude among 15 AudioMoth recorders before (top) and after (bottom) applying Step-3. The boxplots illustrate the reduced variability and increased consistency among the different AudioMoth recorders of the same brand. The amplitude range displayed in the figure is 25 dB.

### 3.3 Data analysis

Figure 7 shows the PDFs of the EAI for each recorder after applying the proposed methodology. Step-1 corrects the PDF mean differences in ACIft and M. The difference in PDFs for ADI, AEI, NP, and H was reduced in Step-2 by limiting the bandwidth to 12kHz, a frequency window in which the recorders have a comparable frequency response, as determined by the coherence measure. The PDFs of the AudioMoth recorder for H, ADI, and AEI remained unchanged after filtering, indicating that information above 12kHz is not notably contributing to these indices. However, AEI would require a more restrictive filter to match both recorders, and the filter neglected all peaks over 12kHz in NP. These undesired effects must be considered in the study design. Finally, Step-3 mainly impacted M and H, which are more sensitive to gain differences (as detailed in Appendix B.4 and Appendix B.5).

**Figure 7.**
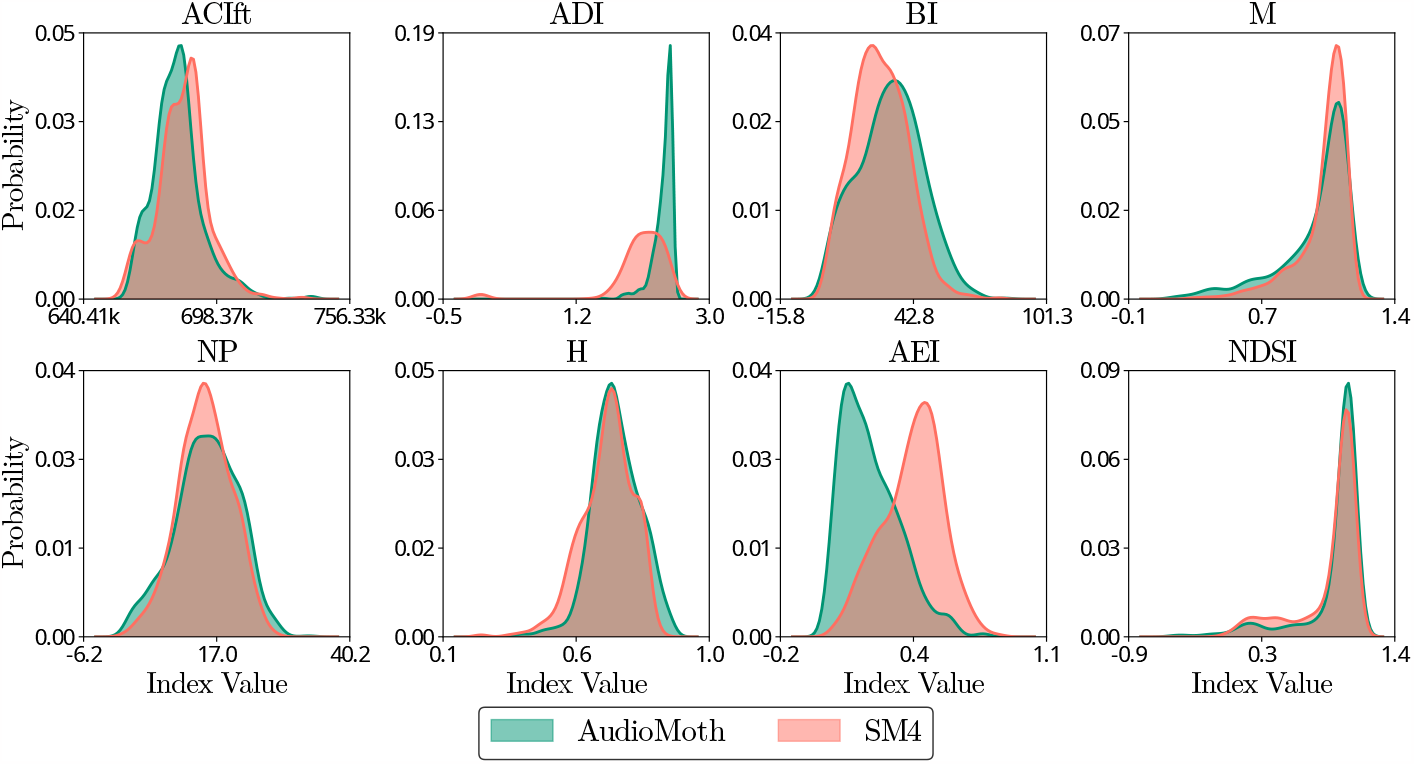
PDFs after applying the proposed recommendations in the EAI calculation using recorders of different characteristics for site S1.

In table 3, we present the KLD and K-S distance metrics calculated for raw and pre-processed data for all indices. The KLD decreases for almost all EAI at the four sites (except NDSI), meaning that the indices are more similar between devices after the pre-processing. The K-S distance is consistent with these results, with reductions of up to 70% in dissimilarity. As expected, the NDSI index did not change after the pre-processing because it is a normalized relation between two frequency bands.

**Table 3:**
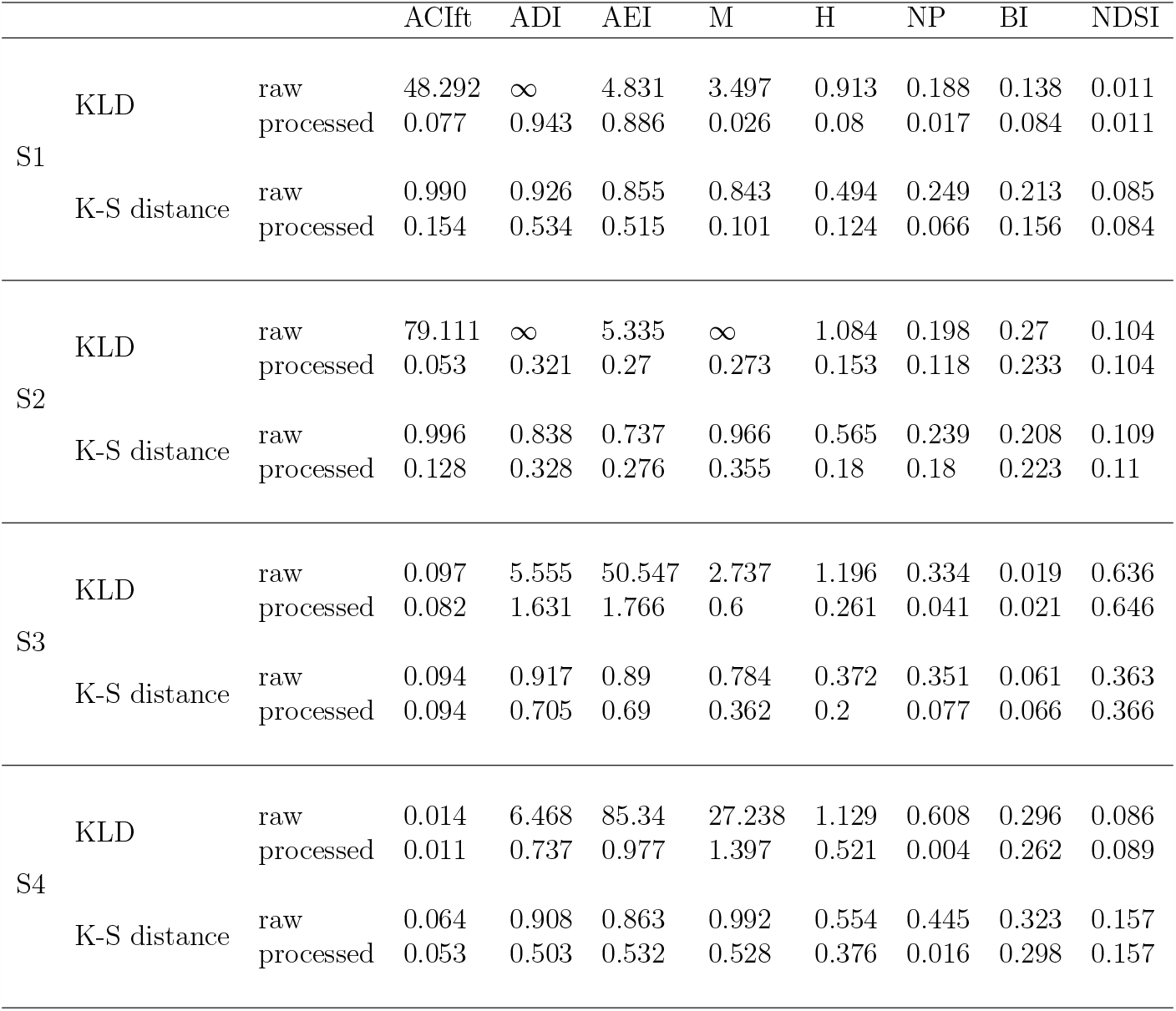
Metrics comparison between EAI using recorders of different characteristics for four geographical locations.

### 3.4 Cross-site/recorder comparison

As a proof-of-principle example, we compared sites S1 and S2 before and after applying the proposed pipeline. Both sites are expected to show similar EAI values. Figure 8 displays the temporal patterns of four selected EAI: (ACIft, ADI, M, and NDSI). The left panels show how the raw indices differ between brands but not intra-brand (except NDSI, as theoretically expected).

**Figure 8.**
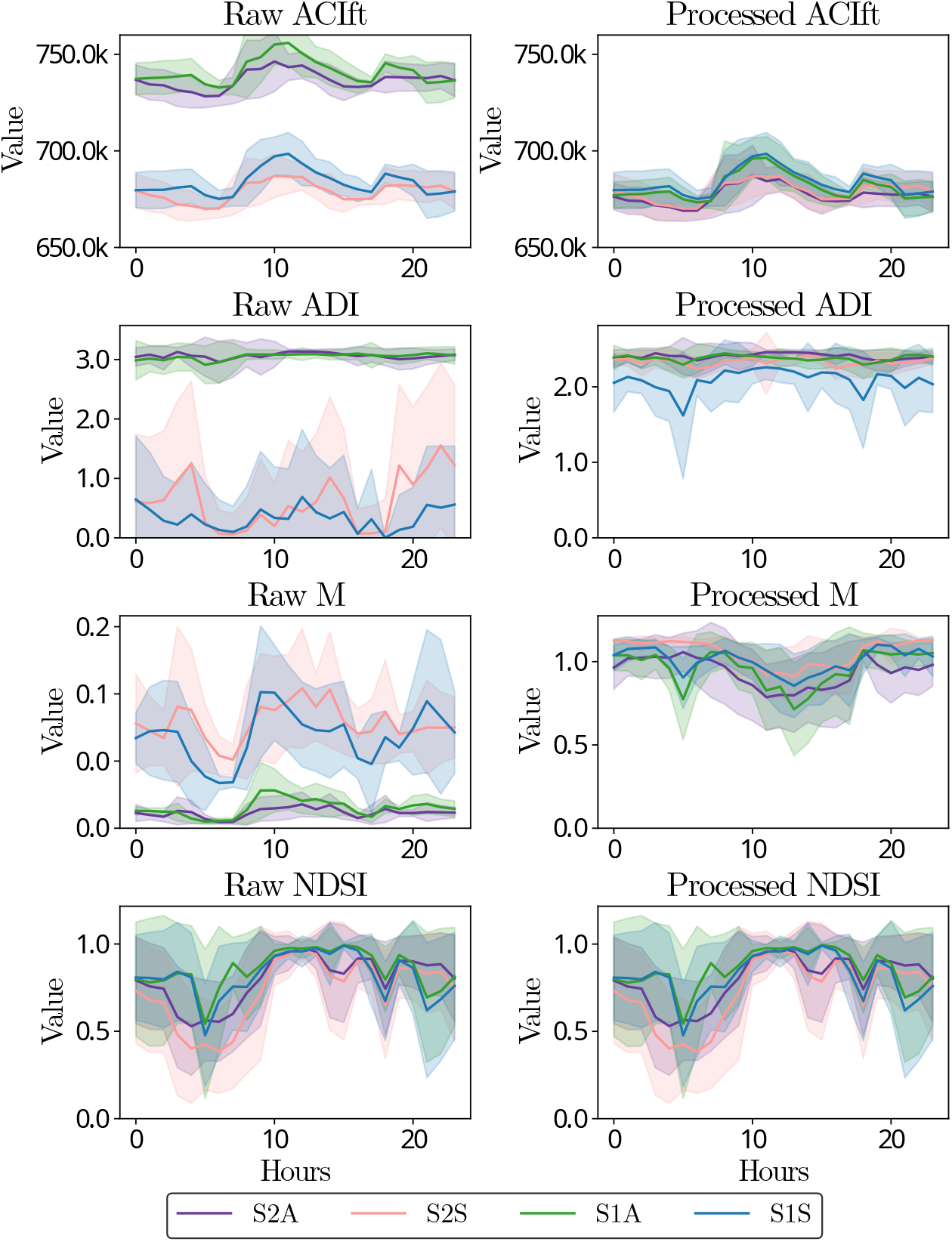
Temporal patterns of EAI showing the effect of the proposed methodology. ACIft was affected by the sampling rate, ADIwas affected by FS variations, M was affected by the signal amplitude, and NDSIremained unchanged as it is robust to these issues. The shadow of each of the lines represents the standard deviation of the index.

Then, the right panels show how the methodology effectively reduces these differences. The other four EAI are shown in Appendix F.

## 4. Discussion

This study investigated the impact of using multiple recorders (even from different brands) over EAI computation. We identified three primary sources of intra- and inter-recorder variability: (i) sampling frequency, (ii) frequency response, and (iii) signal amplitude. To address these issues, we proposed a preprocessing methodology consisting of three steps: resampling, frequency range matching between devices, and normalization. The proposed approach was evaluated using audio recordings simultaneously obtained with AudioMoth and SM4, and our results showed that it effectively reduces up to 70% of device-related biases (in terms of K-S distance). Below, we individually address these sources of variability and their respective proposed solution.

### 4.1 Step-1: Signal resampling

Sampling frequency differences are often disregarded, Adams et al. (2012); Rempel et al. (2013). For example, SM4 recorders commonly employ a sampling rate of 44.1 kHz for audible ranges, while AudioMoth recorders do not allow configuring this sampling rate, being 48 kHz the closest possible value. Our study shows that the behavior of EAI such as ACIft and M is affected by a scale bias when using different sampling rates, preventing a direct comparison among devices. To solve this issue, we employed the Kaiser resampling method proposed in (Liu et al., 2020) to equalize the sampling rates of the devices and we found that effectively decreased the recorder bias for ACIft and M.

Possible problems with resampling methods are that they reduce the spectrum bandwidth, leading to loss of information, particularly in high-frequency bands and that may also deform the signal shape, yielding decreased temporal resolution. Therefore, before applying such correction processes, the effects of subsampling on signal quality must be considered. Future work should include a detailed analysis of this distortion effect and the application of advanced sub-sampling methodologies to minimize these problems.

### 4.2 Step-2: Selecting comparable frequency ranges

The frequency response is a cause of variability between recorders due to the non-linearity they could exhibit (Abrahams et al., 2023). Our theoretical analysis showed that indices with fixed thresholds on gain (ADI and AEI) or that evaluate full spectra (H and NP) are sensitive to this issue, as was effectively observed in our raw data tests. Previous studies have evidenced that differences in the frequency response leads to conflicting conclusions when using indices as proxies for biodiversity or soundscape characteristics (Bradfer-Lawrence et al., 2019b). A common practice to tackle this issue is eliminating microphone noise and other effects caused by the frequency response by applying a high-pass filter with a cut-off frequency of 500 Hz.

However, this strategy does not consider differences between devices for frequencies above 500 Hz, so the variability due as frequency response persists Pieretti et al. (2015).

In this work, we propose to use the coherence measure to statistically compare the frequency bands of the devices and to limit the EAI comparison to those bands where the devices are comparable. Limiting the frequency bands for calculating EAI has already been used to improve the reliability of these indices (Metcalf et al., 2021b). However, the approach of Metcalf et al. (2021b) requires prior knowledge of the soundscape and the masking of signals to determine the frequency ranges to be selected; i.e., an expert would require to analyze the areas beforehand, which is a disadvantage compared to our proposed automatic approach.

Our results showed that the coherence measure provides a consistent approach to identify potential discrepancies in frequency responses when comparing recorders. Nevertheless, limiting the frequency ranges to calculate EAI neglects potentially relevant biological information. As future work, identifying signal processing techniques that correct this behavior without ruling out information from these frequency bands is required.

### 4.3 Step-3: Normalize the signal amplitude

The signal amplitude refers to the audio intensity at a given moment, which is affected by different factors in the acquisition process, such as The microphone frequency response, compression format, and coding (sampling depth) (Browning et al., 2017). When comparing audio acquired with different devices, it was found that the amplitude of audio signals will likely be different (even with same-brand recorders). Despite the importance of this issue, many works focus on the behavioral pattern of the index without considering its magnitude (Tuneu-Corral et al., 2020), losing reproducibility among studies. Our theoretical analysis showed that this amplitude variation creates biases in those EAI calculated in the time domain that work with absolute values (and do not normalize), such as M and H.

To solve this issue, we normalized the signal with its RMS value before calculating the index to ensure that they have the same reference value. Furthermore, RMS normalization does not limit the audio to a specific range, thus avoiding using other signal normalization techniques that could suppress relevant information (Vu et al., 2018). As a result, variabilities in M and H were reduced between devices. This improvement is less evident in H at site S4 (see Table 3). It may be due to the complexity of the sound space at this location, since the initial PDFs have a different shape from other locations. Finally, when this normalization is used, it should be considered that it modifies the EAI scale, which could be an inconvenience if a direct interpretation of its magnitude is required.

Contrasted with the studies elucidated in the literature review depicted in Table Appendix A, our full methodology distinguishes itself through its all-encompassing approach to rectifying the bias introduced by recording devices in acoustic indices. As gleaned from the literature, intra-brand evaluations predominantly concentrate on diagnosing the issue instead of proposing an effective remedy to counteract the bias stemming from recording devices (Adams et al., 2012; Rempel et al., 2013; Péerez-Granados et al., 2019). Furthermore, certain studies delve into the variability of acoustic indices, often emphasizing filtering techniques without specifying precise frequency ranges or protocol modifications to alleviate the bias (Metcalf et al., 2021b; Cifuentes et al., 2021; Hyland et al., 2023a). In contrast, our methodology encompasses a broader spectrum by incorporating these solutions more comprehensively. Through this encompassing perspective, our approach equips researchers with a powerful tool to mitigate variability in ecoacoustic indices, enabling more dependable and comparable outcomes across diverse recording scenarios.

### 4.4 Proof-of-principle example

Our proof-of-principle example demonstrates how our full pipeline allows comparing across sites with all EAI despite using devices from different brands. EAI are not commonly used for comparing sites because they have shown contradictory correlations with biological variables depending on the study site (Bradfer-Lawrence et al., 2019a). Nevertheless, to use our methodology breaks this limitation and opens the ecoacoustics field to new kinds of analyses. Moreover, by making EAI comparable across studies, we can use them in new applications such as machine learning and deep learning, which require features in the same space, i.e., without considering the device with which they were acquired (Purwins et al., 2019).

Note that our initial assumption of similar behavior between sites S1 and S2 was solely based on the opinion of the biologist who placed the recorders. However, our results align with this assumption and demonstrate similar biological characteristics between these two locations. Nonetheless, this is only a limited proof of principle, and further analyses should be carried out to validate this assumption.

## 5. Conclusion

In this study, we have demonstrated that the recording device introduces an understudied source of variability, thereby influencing EAI and potentially affecting the conclusions and reproducibility of soundscape monitoring studies. We identified three issues related to using multiple recorders: differences in sampling frequency, variations in microphone frequency response, and disparities in gain settings. Our proposed signal pre-processing pipeline, designed for easy implementation in three steps, effectively mitigates recorder-induced bias in EAI and allows directly comparing them across diverse audio recording devices. Our results unveil a remarkable reduction of up to 70% in biases attributed to recording devices (measured by K-S distance). This pipeline will be a starting point for developing strategies to perform multi-recorder analyses.

## 6. Acknowledgments

This work was supported by Universidad de Antioquia, Instituto Tecnologico Metropolitano de Medellín, Alexander von Humboldt Institute for Research on Biological Resources, and The Colombian National Fund for Science, Technology and Innovation, Francisco Jose de Caldas - MINCIENCIAS (Colombia). [Program No. 111585269779].

## Appendix A. Literature review on variability in ecoacoustic Data

**Table A.4:** Comprehensive review of literature on strategies addressing variability in ecoacoustic data: reduction and source identification approaches.

| Paper | Recorder | Brand | Acoustic Indices | Source variability | Proposal solution |
| --- | --- | --- | --- | --- | --- |
| Bradfer-Lawrence et al. (2023) |  |  | ✓ | - Quantity of recordings<br>- Recording schedules<br>- Microphone noise | - Collecting high quality recording<br>- Collecting sufficient recordings<br>- Avoid temporal subsampling<br>- Design the sampling protocol<br>- Interpreting indices considering sources of variability |
| Hyland et al. (2023b) | Single | Intra | ✓ | - Environmental conditions<br>- Noise sources | - Filters at different frequencies<br>- habitat-dependent frequency ranges |
| Cifuentes et al. (2021) | Single | Intra | ✓ | - Sampling protocol<br>- Length of recordings | - Design the sampling protocol considering sample duration and effect on indices |
| Metcalfe et al. (2021b) | Multiple | Intra | ✓ | - Signal masking<br>- Noise source | - Calculate the indices in specific time and frequency ranges depending on the sound-space of the habitat |
| Bradfer-Lawrence et al. (2019a) | Multiple | Intra | ✓ | - Quantity of recordings<br>- Recording schedules<br>- Microphone noise | - High-pass filtering at 500 Hz<br>- Collect 120 hr of audio recordings per site<br>- Avoid temporal subsampling<br>- Consider more than just a single portion of the data |
| Pérez-Granados et al. (2019) | Multiple | Inter |  | - Recording device<br>- Recorder distance | - Explore various recorders and an optimal selection for specific applications |
| Fairbrass et al. (2017) | Single | Intra | ✓ | - Abiotic noise | - N/A |
| Gasc et al. (2015) | Simulated |  | ✓ | - Sound composition<br>- Signals masking | - Interpreting indices considering sources of variability |
| Rempel et al. (2013) | Multiple | Inter | - | - Recording device<br>- Internal noise of the recorder<br>- Microphone frequency response | - N/A |
| Adams et al. (2012) | Multiple | Inter |  | - Recording device | - Considering the influence of recorders in sampling design |

## Appendix B. Ecological acoustic indices

### Appendix B.1. Acoustic complexity index (ACIft)

The Acoustic Complexity Index (ACIft) was proposed by Pieretti et al. (2011). It is based on the biological hypothesis that many biotic sounds have intrinsic variability in their frequency bands, in contrast to anthropophonic noise that tends to be constant, such as car and airplane traffic. It was initially proposed to quantify bird calls since they present significant intensity variations in the vocalization frequencies. However, its use has been extended to different tasks, such as analyzing habitat quality, communities, and the richness of various species Buxton et al. (2018).

ACIft index works in the time-frequency domain, and its analytical approach is based on the spectrogram matrix *S*. Fourier coefficients compound this matrix and encode the signal intensity in time and frequency bins. It is obtained by first computing the Fast Fourier Transform (FFT) (Nussbaumer, 1981) of the audio signal and then calculating the absolute difference of intensities in the adjacent bins of the time axis. This operation can be interpreted as the time derivative of *S*.

The next step is to normalize each row of the matrix obtained by deriving *S* over the sum of intensities in the frequency band. This provides relative values and reduces the effect of the call distance on the microphone. Finally, the intensity differences in time and frequency are summed, as shown in Equation B.1:

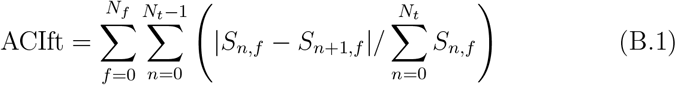

where *N*_*f*_ and *N*_*t*_ are the number of time and frequency bins, respectively, and *S*_*n,f*_ is the bin at the *n*-th time sample and *f* -th frequency sample.

The ACIft is robust to using different recorders with different gains (e.g., due to different brands of microphones) because it is based on relative values. However, it is sensitive to different sample frequencies since the number of bins in *S* depends on the sampling frequency and would cause a bias in (B.1). Finally, this index is sensitive to recordings with different geophonic sources (such as rain or wind) or non-constant sources of anthropophony (e.g., planes and cars) because they increase the intensity differences and falsely increase the biological complexity.

### Appendix B.2. Acoustic diversity index (ADI)

The Acoustic Diversity Index (ADI) was proposed by Villanueva-Rivera et al. (2011) to determine the occupancy degree of acoustic niches. It is calculated from the spectrogram of the signal in decibel scale partitioned into frequency bands of 1kHz bandwidth, after which the spectrogram is binarized using a threshold (−50dBFS by default) to eliminate background noise. With the binary matrix *Z ∈ N*_*t*_ *× N*_*i*_, the probability of occupancy at each frequency band *ρ*_*i*_ is calculated as shown in (B.2) for each *i* = 1, …, *N*_*ı*_frequency band.

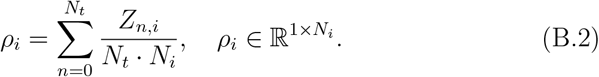

Finally, the ADI is calculated based on Shannon entropy of the occupancy probability:

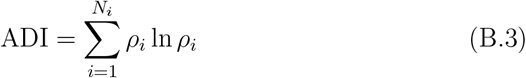

Opposite to ACIft, ADI is sensitive to recorders with different gains because the binarization threshold would vary among devices. As this index works with frequency bands, the frequency response of the microphone is critical because one can, for example, determine in which frequency bands the gains of different devices are comparable. Otherwise, the occupancy level of acoustic niches would not be comparable. On the other hand, ADI is robust against sampling frequency variations because the frequency domain is collapsed. Finally, it is only sensitive to artifacts (e.g., geophony or anthropophony) with energy content in several bands (such as rain).

### Appendix B.3. Acoustic evenness index (AEI)

The Acoustic Evenness Index (AEI) is calculated following the same procedure explained in the ADI index to obtain the probability distribution function (see (B.2)), but it then continues with the Gini coefficient Gini (1912)

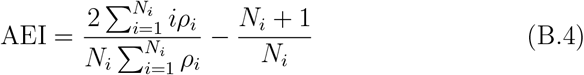

computation: which measures the inequality between the values of the probability distribution. Ecologists use this index to estimate the uniformity of species in different acoustic niches Villanueva-Rivera et al. (2011). Additionally, the AEI index has been used to differentiate acoustic activity at different times of the day and in different habitats Droge et al. (2021).

Its main difference with ADI is that instead of using a logarithmic weighting (see (B.3)), it uses a pseudo-square weighting (see (B.4)). This difference results in a negative correlation between AEI and ADI indices, as demonstrated by (Peet, 1974). Therefore, they require different interpretations, which makes sense as the highest diversity reduces uniformity and vice-versa. However, since these indices may provide similar information, using both indices may sometimes be redundant (Eldridge et al., 2018). Because of their similarities, they have the same disadvantages against artifacts and the use of different recorders.

### Appendix B.4. Median of the amplitude (M)

The amplitude index M was proposed by Depraetere et al. (2012) as an indicator of the number of animal calls under the hypothesis that the more calls, the higher the signal amplitude peaks, which are obtained with the median signal envelope:

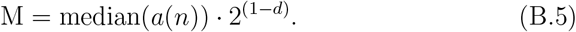

This envelope *a*(*n*) 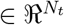 is commonly computed using the Hilbert transform (Johansson, 1999), which provides a smoother signal and encodes its amplitude at all time intervals, as shown in Figure B.9. Where *n* is the number of temporal samples of the envelope. Note that the sampling frequency of the envelope is lower than that of the original signal. The depth of bits *d* indicates the resolution at which the audio can be acquired, which defines the dynamic range of amplitude. Each bit corresponds to 6 dB, meaning that a recording with a 16 bits depth can capture a dynamic range of 96 dB. Any amplitude outside this range will not be captured or will be distorted. The signal envelope is scaled by a factor of the digitization depth of the recording to avoid biases. As a final step, the authors recommend a second normalization of the M index to a range of [0,1] by dividing it by its maximum value to avoid too small values (of order 10^−3^) and facilitate interpretation.

The main disadvantages of M are that several factors, such as the microphone distance to the transmitter and the habitat structure, may influence the signal amplitude and, therefore, the envelope. These factors make it sensitive to environments with high-intensity noise sources, causing a high richness of false calls. This problem is notably when using different recorders because M cannot account for recordings obtained from microphones with different gains. Consequently, even with its double normalization, the problem of non-comparability among recordings obtained from different devices persists. Additionally, M is also sensitive to recordings with different sampling frequencies, as the smoothing of the Hilbert transform is related to the number of samples (although this is usually a minor problem in practice). Thus, when using this index with different devices, it is essential to ensure that the audio recordings have the same sampling frequency to avoid potential errors.

**Figure B.9:**
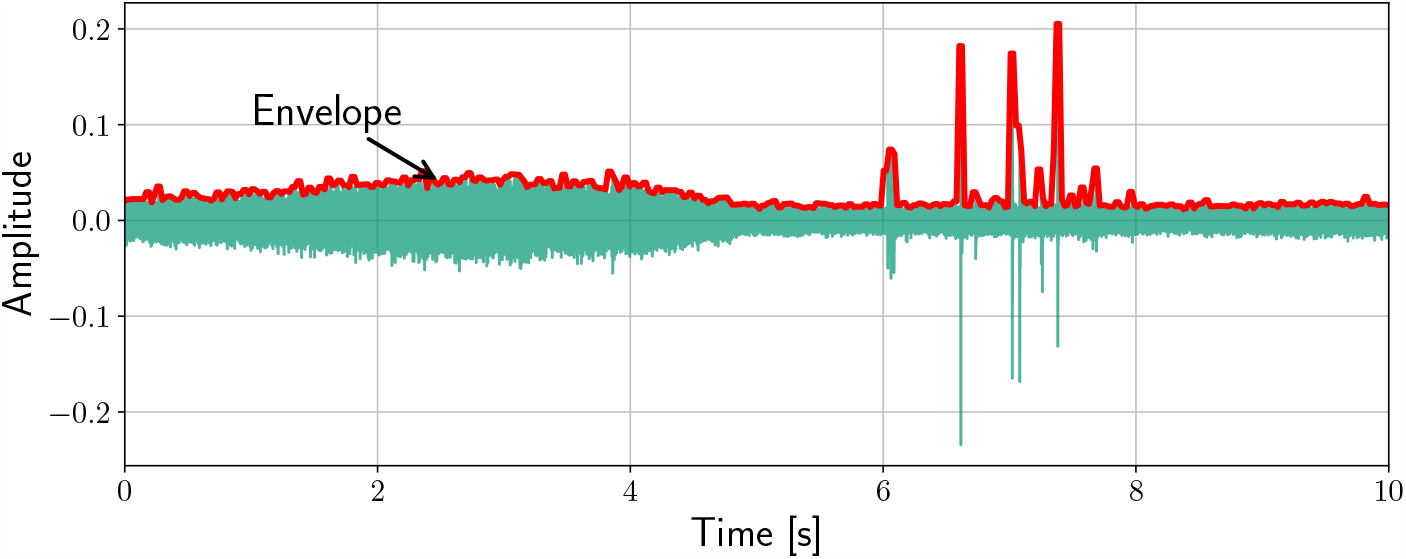
Envelope of the audio signal used to compute the acoustic index M.

### Appendix B.5. Acoustic Entropy index (H)

The Entropy Index (H) was proposed by Sueur et al. (2008) as an indicator of heterogeneity and is based on estimating the Shannon uniformity over time and frequency. First, the temporal entropy index *H*_*t*_ is computed by normalizing the signal envelope *a*(*n*) over time:

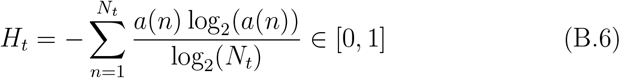

Each time instant *N*_*t*_ has an associated amplitude value, normalized by dividing it by *log*_2_(*N*_*t*_), corresponding to the maximum value of the Shannon index and depending on the number of amplitude samples (or categories in information theory). The main advantage of using a base two logarithm is its sensitivity to infrequent categories. Hence, low probability categories are influential in acoustic diversity. Furthermore, normalization provides a relative value that decreases bias from constant noise sources.

The second part corresponds to the spectral entropy index *H*_*f*_, which measures the Shannon uniformity of the frequency spectrum. *H*_*f*_ is calculated using the spectral probability mass function *s*(*f*), obtained by averaging the spectrogram on the time axis and normalizing it over the number of frequency bins *N*_*f*_. Therefore,

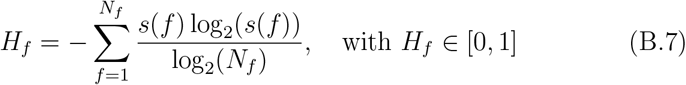

The index is calculated as the multiplication of the temporal and spectral entropy indices: H = *H*_*t*_ · *H*_*f*_. Its purpose is to encode the acoustic complexity by entropy, assuming that it increases with species richness and the number of individuals. In this case, complexity is a synonym for heterogeneity, and many EAI are based on this same hypothesis.

The way that *H*_*t*_ is computed reduces its sensitivity to gain differences among devices, as well as different sampling frequencies. However, it is sensitive to broadband noise sources (such as traffic, rain, and wind, among others). On the other hand, the normalization term in *H*_*f*_ also reduces its dependency on the frequency sample, but again broadband noise sources affect it.

### Appendix B.6. Number of peaks (NP)

The Number of Peaks index (NP) was proposed by Gasc et al. (2013) as an indicator of acoustic complexity using the number of prominent peaks in frequency (see Figure B.10). The number of peaks is assumed as a spectral profile of the acoustic richness of the habitat. The NP index is calculated using the average spectrum obtained with *H*_*f*_ but scaled to [0, 1]. Subsequently, traditional peak detection algorithms (such as proposed by Dumpala et al. (1982) or Sueur (2018)) are used to find prominent peaks. In practice, only peaks with slopes greater than 0.01 amplitude units and separated by at least 200 Hz are kept.

The NP index is based on the assumption that the number of peaks in frequency is related to the acoustic activity in different niches. Its main advantage is that it is less sensitive to environment noise than most EAI because the noise sources tend to be limited to a specific frequency band or have pseudo-constant behavior among frequency bands. It is also robust to gain and sample frequency differences among devices because of the normalization in amplitude and the fact that it does not perform computations over frequency bins.

**Figure B.10:**
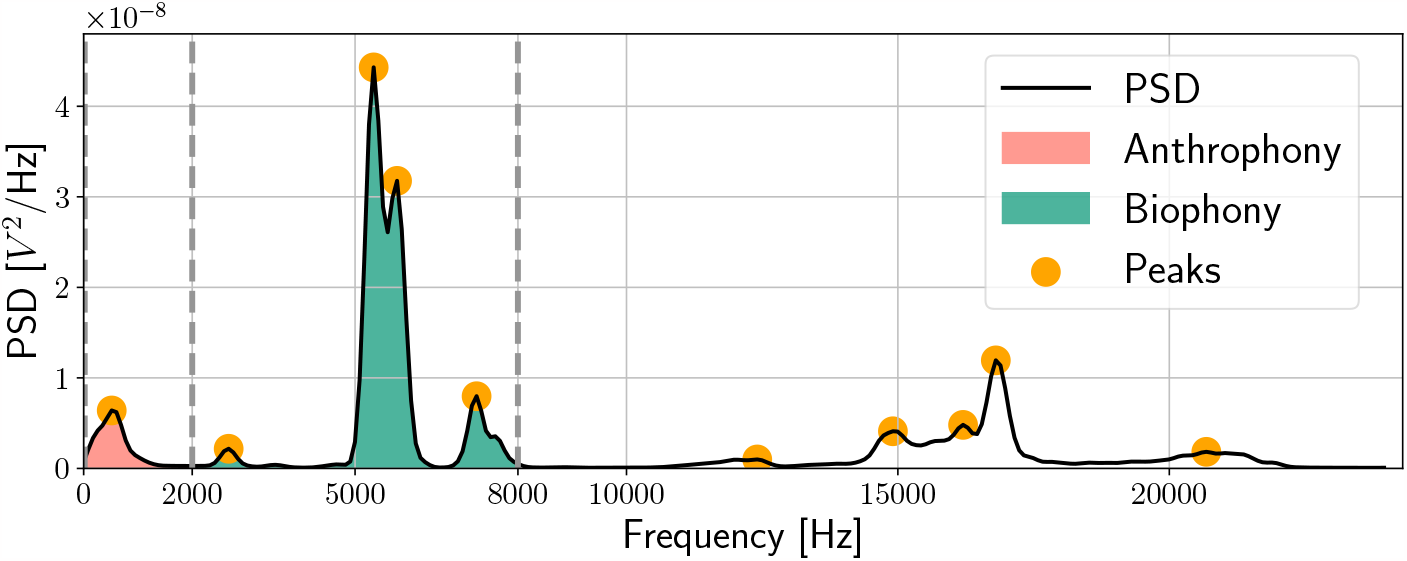
Defined frequency ranges for biophony, anthrophony, and spectrum peaks.

### Appendix B.7. Bioacoustics index (BI)

The bioacoustic index (BI) was initially proposed by Boelman et al. (2007) to assess the relative abundance of birds in the biophony frequency bands, which range from 2 kHz to 8 kHz. However, it has been used to estimate the abundance of different species by expanding the range of the frequency bands defined for biophony, such as [2 kHz–10 kHz] or [2 kHz–12 kHz], depending on the species being studied (Ferreira et al., 2018). The BI index uses the spectrum mean in the same way as *H*_*f*_ but in the decibel scale. Then, the area under the curve is evaluated in the limits defined for biophony:

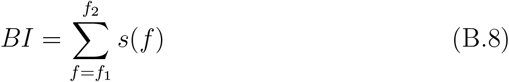

which in the original approach of the index correspond to *f*_1_ = 2 kHz and *f*_2_ = 8 kHz (see Figure B.10).

BI is susceptible to broadband noise sources that overlap with the biophony band. Additionally, both the sampling frequency of the recorder and the frequency response of the microphone in the biophony band can strongly alter the results when using different devices because of the lack of normalization in the number of spectrum partitions and the device gain will notably bias the sum in (B.8).

### Appendix B.8. Normalized difference soundspace (NDSI)

The Normalized Difference Soundscape Index (NDSI) was proposed by Kasten et al. (2012) to estimate the level of anthropogenic disturbance in the soundscape. It is defined by a ratio of human-generated acoustic component anthrophony *α* and a biological component (biophony) given by the bioacoustics index BI. The anthrophony is calculated using the same procedure to compute BI, but the limits of the frequency bands are changed to *f*_1_ = 1 kHz and *f*_2_ = 2 kHz (see Figure B.10). Then, the NDSI index is computed as follows:

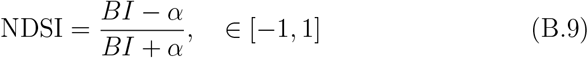

The NDSI index is considered an indicator of the distribution of the soundscape at different times of the day. The NDSI is bounded between [−1, 1], where NDSI = 1 indicates that the signal does not contain anthrophony. However, some species that make calls at low frequencies can generate values of NDSI < 0.8, so there is an acceptable variability considering the high complexity of biological systems. On the contrary, a NDSI = −1 indicates no biophony in the recording. The high interpretability of NDSI has led it to be one of the most widely used indices in soundscape studies. However, as it comes from BI, all its weaknesses are translated to NDSI. Additionally, the algorithm may fail when there is a noise below 500Hz (to avoid it, some authors recommend filtering before computing the NDSI).

## Appendix C. Site: S2

### Appendix C.1. Coherence

Figure C.11 presents the results of applying the coherence measure to the data from site S2. In the top panel, note how in the frequency range of 7-8kHz, the frequency responses are not comparable as they do not exceed the established threshold. This is due to the behavior change exhibited by the SM4 recorder at this frequency range, which can be observed in the bottom panel of Figure C.11. In addition, the frequency response at this site is not comparable over 12kHz, which is consistent with the other sites.

### Appendix C.2. indices

Figure C.12 shows the PSDs of the indices for the two recording devices at site S2. They exhibit similar behavior to that reported for site S1 in Figure 3. However, the BI PDFs show a more notorious difference among devices, contrary to that observed in the other analyzed sites.

Figure C.13 confirms the consistency of the corrections with the results obtained in site S1.

**Figure C.11:**
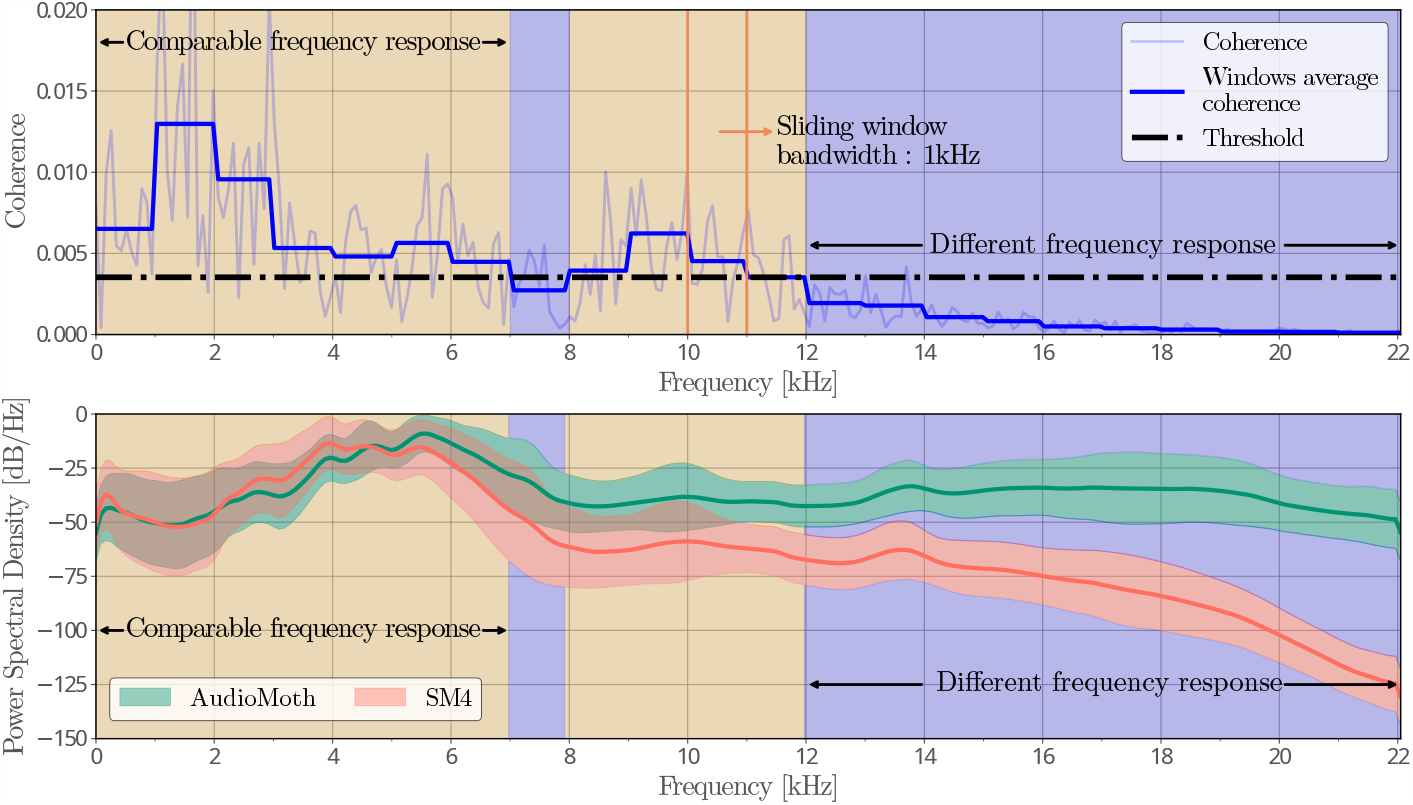
(Top) Coherence sliding window for detecting comparable frequency ranges between recorders in site S2. (Bottom) PSD average of subsampled signals.

**Figure C.12:**
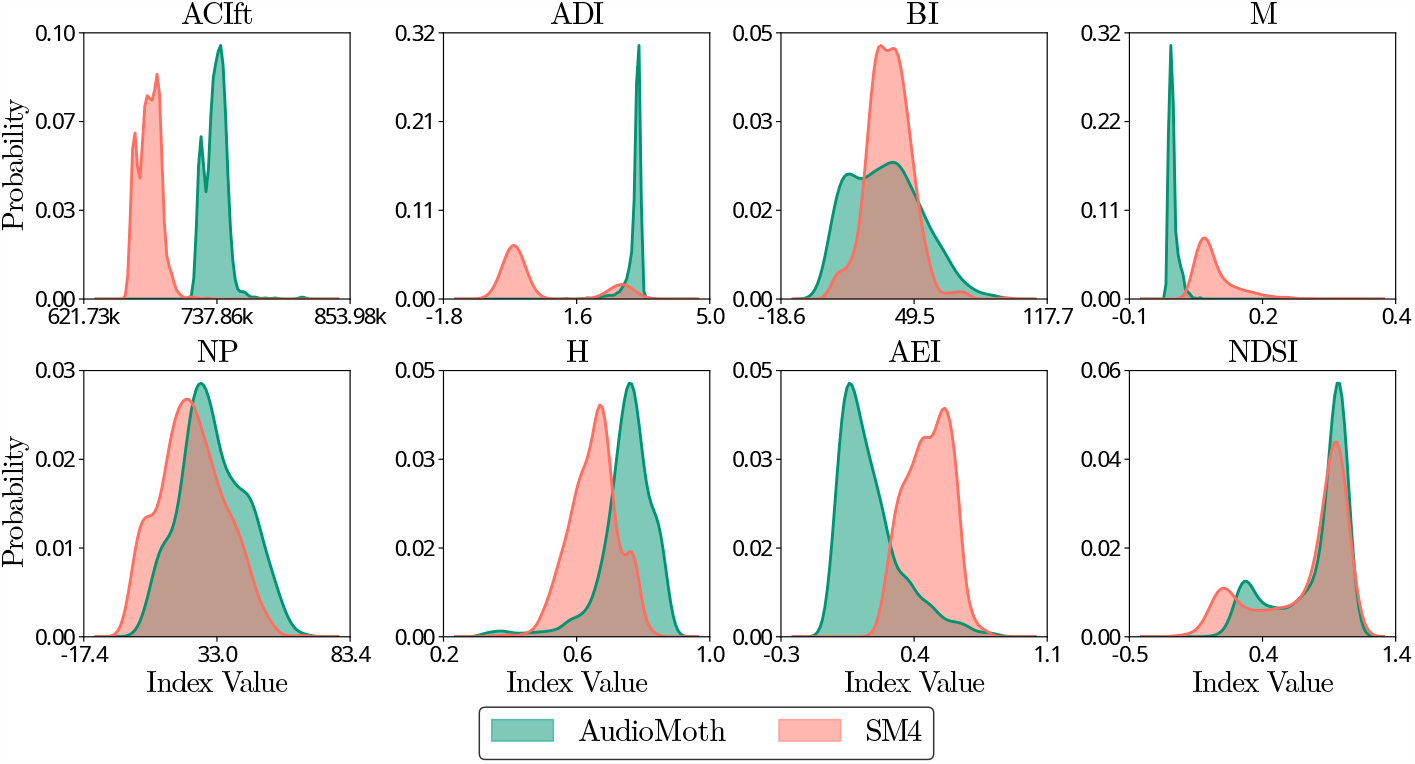
PDFs of the raw EAI calculated using recorders of different characteristics for site S2.

**Figure C.13:**
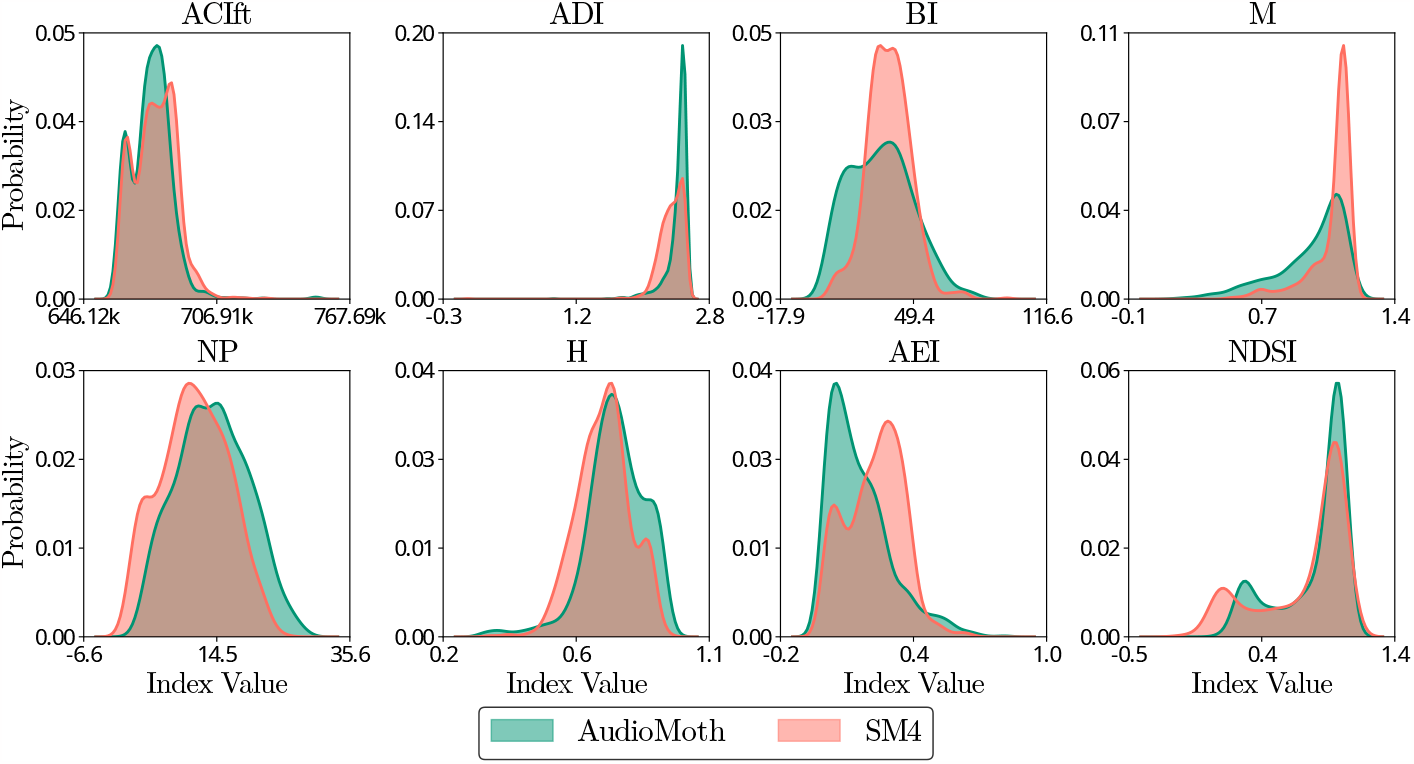
PDFs of the processed EAI calculated using recorders of different characteristics for site S2.

## Appendix D. Site: S3

The behavior of EAI before pre-processing for site S3 is shown in Figure D.14. As expected, the ACIft index does not present any difference in mean between PDFs, as the audio recordings were made with the same sampling rate for both devices. An atypical behavior is observed in NDSI, presenting a higher difference than the other study points. The rest of the indices exhibit a behavior similar to that already reported.

Figure D.15 shows how the difference between PDFs of the devices is reduced when the proposed processing approach is applied. As discussed above, M shows a contrasting behavior in this case, as it slightly increases the dissimilarity between devices when applying the proposed approach.

## Appendix E. Site: S4

Site S4 did not exhibit any atypical behavior in the PDFs of the EAI for the different recording devices. All results are consistent with those reported in the previous sites, as shown in Figure E.16. Similarly, when applying the proposed preprocessing steps, the differences between the devices are consistently reduced, indicating that methodologies are applicable in different sites, showing similar behavior, as seen in Figure E.17.

**Figure D.14:**
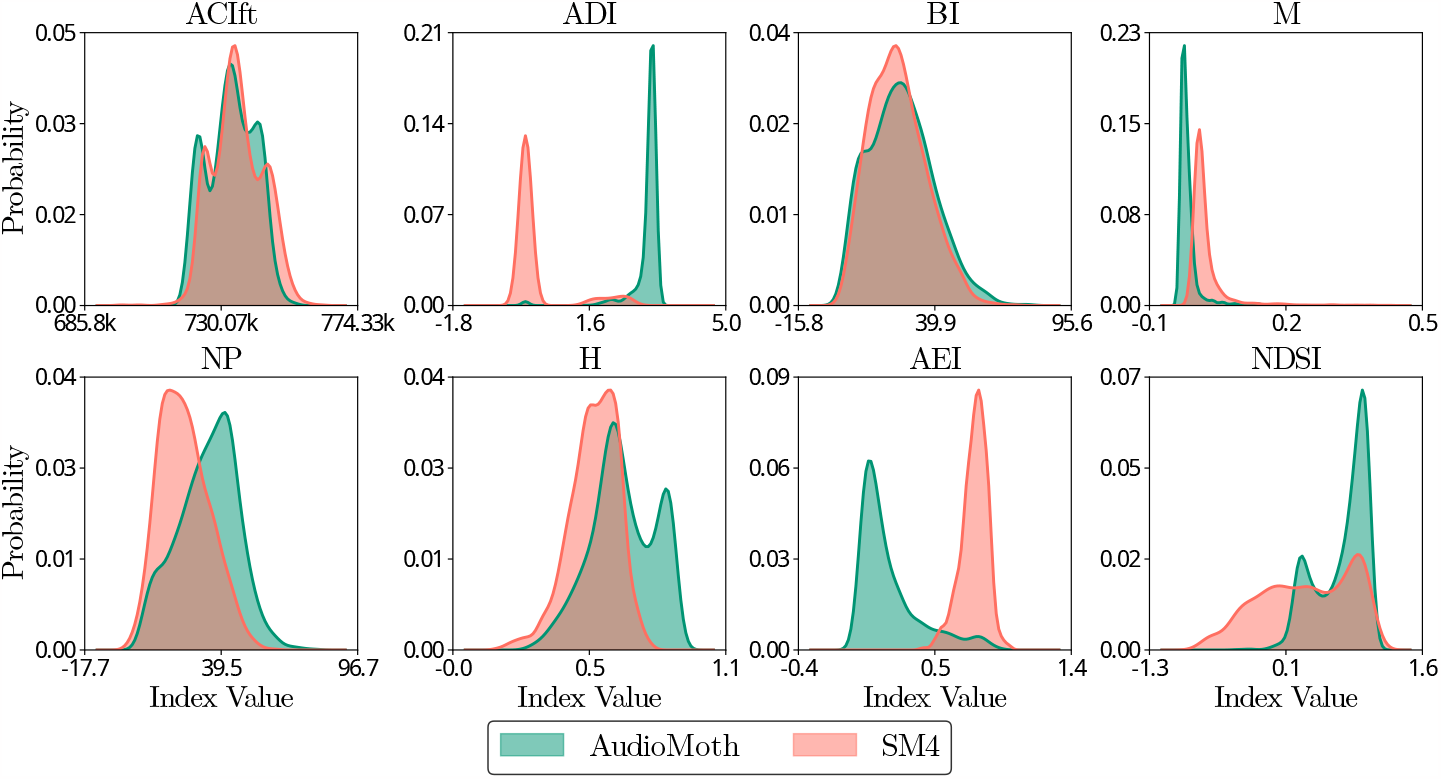
PDFs of the raw EAI calculated using recorders of different characteristics for site S3.

**Figure D.15:**
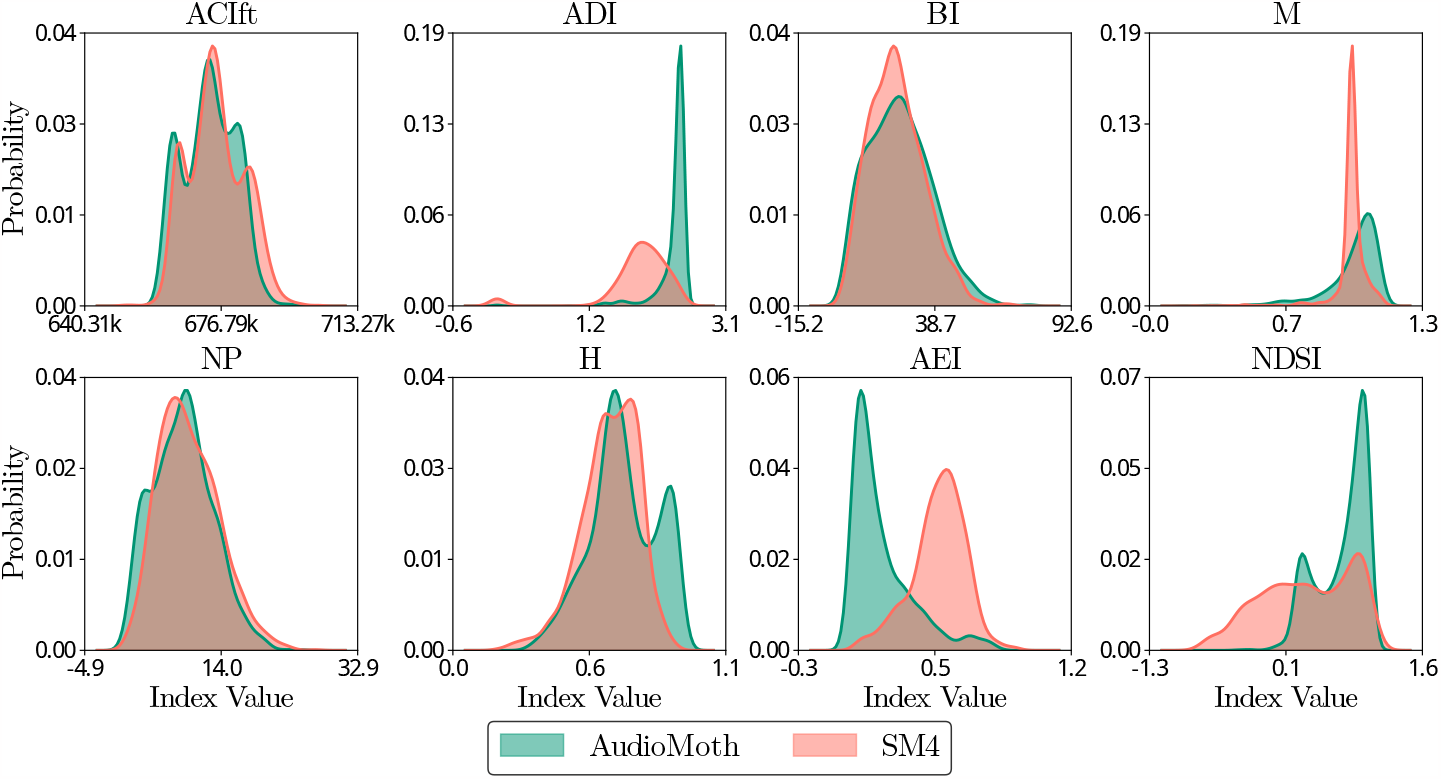
PDFs of the processed EAI calculated using recorders of different characteristics for site S3.

**Figure E.16:**
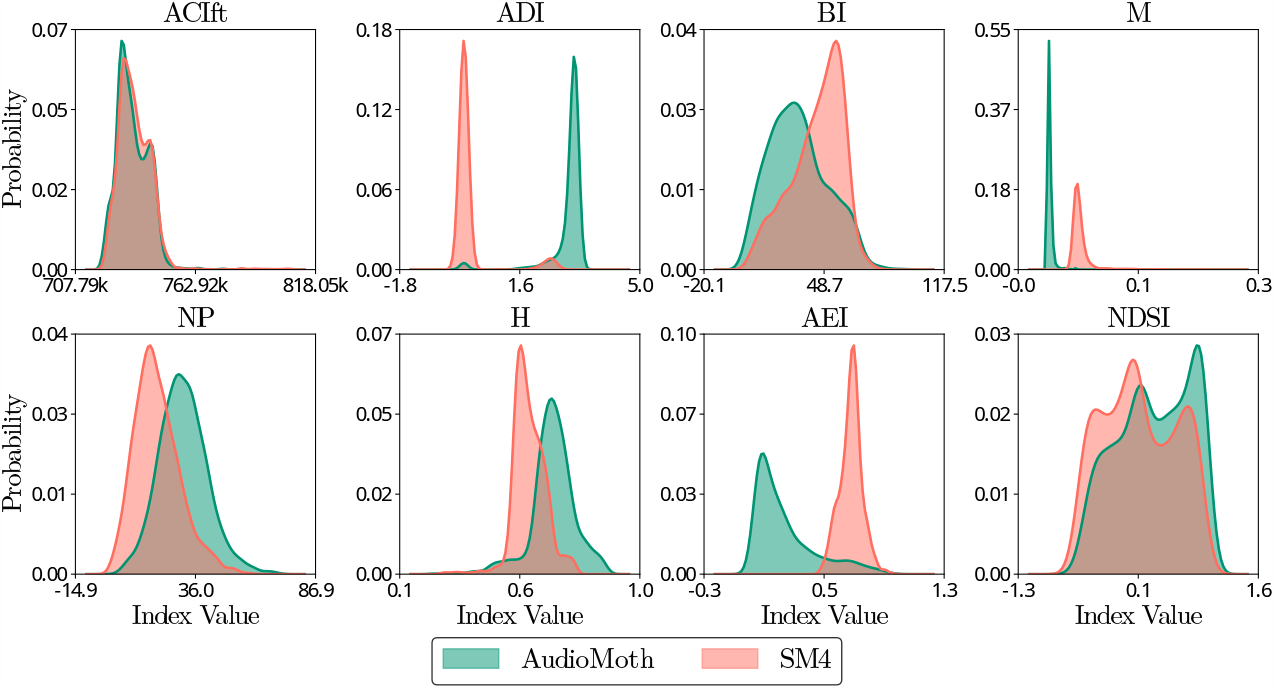
PDFs of the raw EAI calculated using recorders of different characteristics for site S4.

**Figure E.17:**
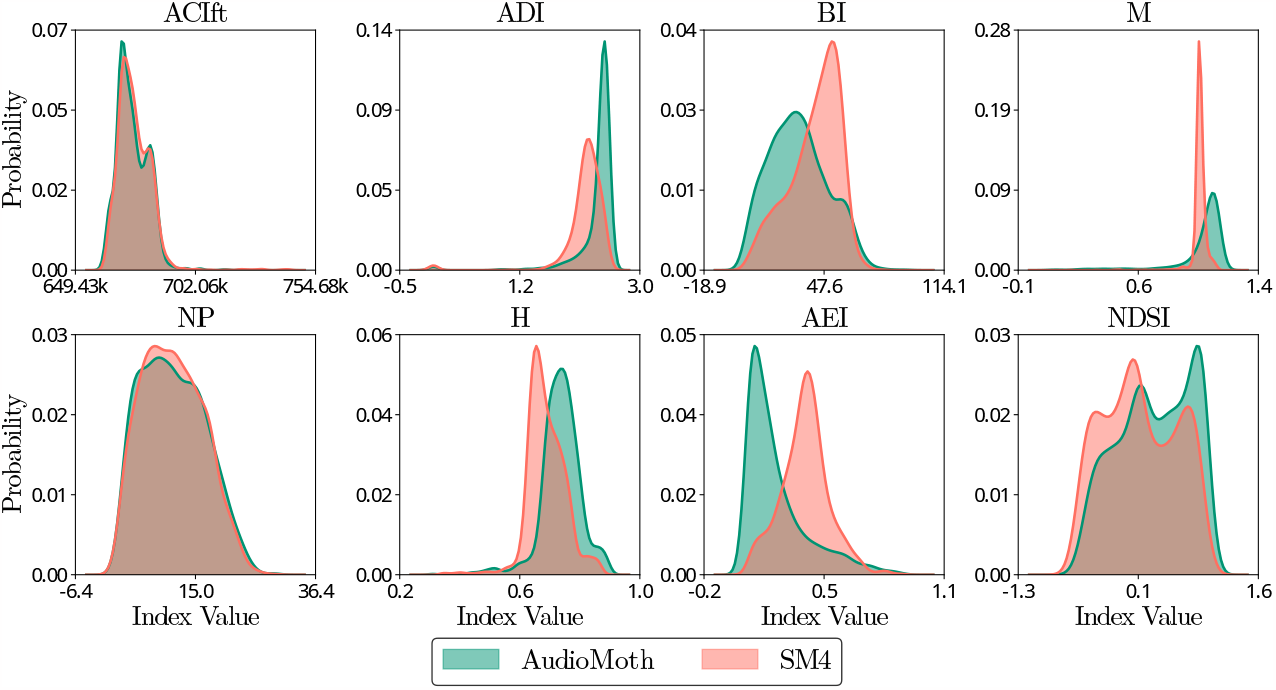
PDFs of the processed EAI calculated using recorders of different characteristics for site S4.

## Appendix F. Site comparison

In this study, we compared the acoustic properties of sites S1 and S2 using the AudioMoth and SM4 recorders at each site. Figure F.18 displays the temporal patterns of AEI, BI, H, and NP indices before and after applying our proposed methodology. Reduced differences within EAI are observed. Notably, AEI showed reduced variability, although there were still noticeable differences in the temporal patterns between the recorders. The other indices did not show such notable differences with raw data, but we still were able to reduce the existing differences through our methodology.

**Figure F.18:**
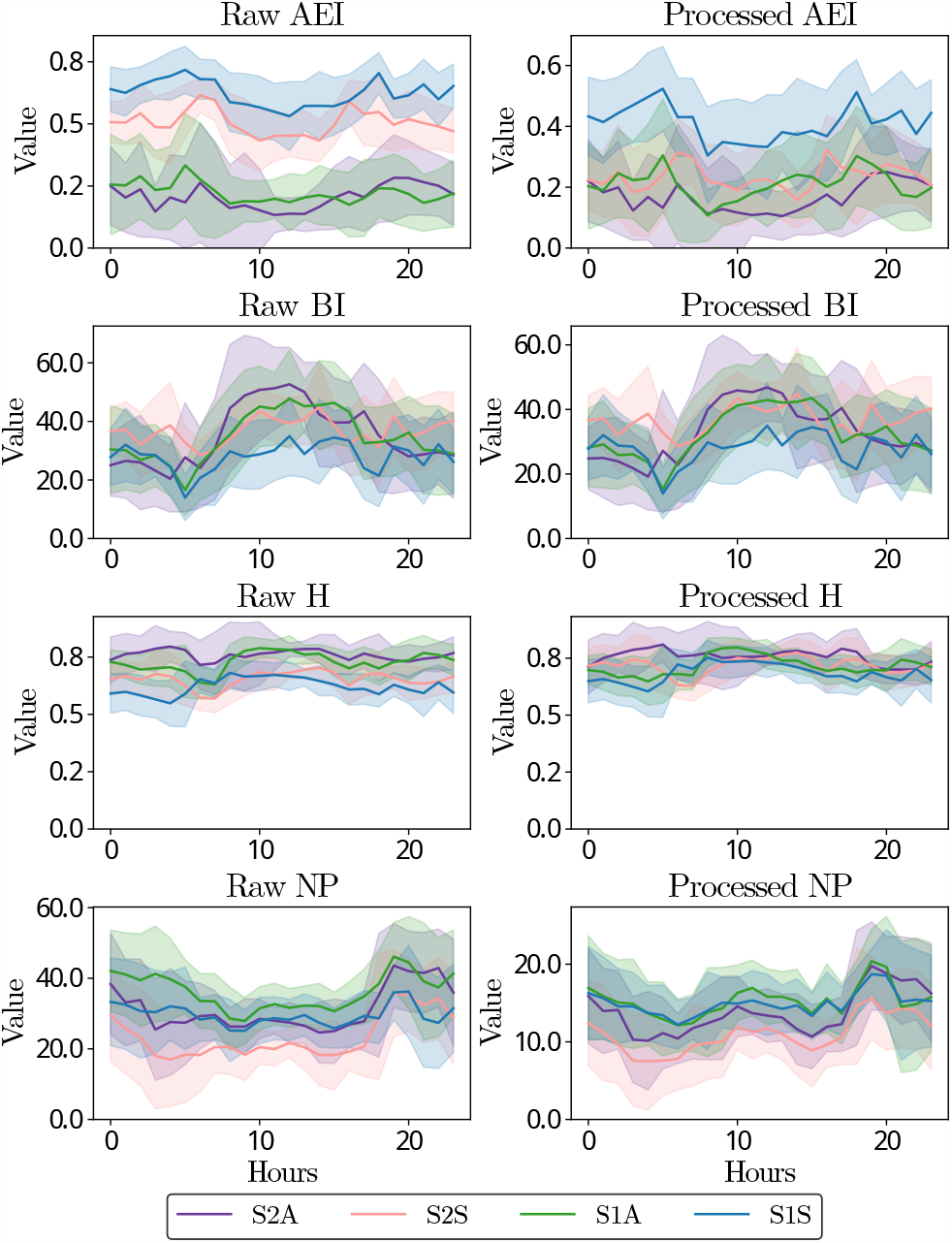
Temporal patterns of EAI showing the effect of the proposed methodology.

